# Dual proteomic signature of immune cells and *Yersinia pestis* upon blood infection

**DOI:** 10.1101/2023.06.19.545537

**Authors:** Pierre Lê-Bury, Thibaut Douché, Quentin Giai Gianetto, Mariette Matondo, Javier Pizarro-Cerdá, Olivier Dussurget

## Abstract

Emerging and reemerging infectious diseases represent major public health concerns. The urgent need for infection control measures requires deep understanding of molecular pathogenesis. Global approaches to study biological systems such as mass-spectrometry based proteomics benefited from groundbreaking physical and bioinformatical technological developments over recent years. However, dual proteomic study of highly pathogenic microorganisms and their hosts in complex matrices encountered during infection remains challenging due to high protein dynamic range of samples and requirements imposed in biosafety level 3 or 4 laboratories. Here, we constructed a dual proteomic pipeline of *Yersinia pestis* in human blood and plasma, mirroring bacteremic phase of plague. We provide the most complete *Y. pestis* proteome revealing a major reshaping of important bacterial pathways such as methionine biosynthesis and iron acquisition in human plasma. Remarkably, proteomic profiling in human blood highlights a greater *Yersinia* outer proteins intoxication of monocytes than neutrophils. Our study unravels global expression changes and points to a specific pathogenic signature during infection, paving the way for future exploration of proteomes in the complex context of host-pathogen interactions.

**Subject Categories:** Microbiology, Virology and Host Pathogen Interaction, Proteomics

## Introduction

Infectious diseases emerging and reemerging worldwide are major public health issues. The understanding of microbial pathogenesis necessary for infection control immensely benefits from technological progress. In particular, the current omics era shows spectacular advances allowing affordable, fast and reliable genome sequencing and deep-coverage of whole-genome transcriptomes. Simultaneously, whole-proteome exploration is quickly gaining ground as the next step for functional studies in systems biology. Classically, proteomics pipeline includes 3 steps: sample preparation, followed by data acquisition in mass-spectrometry and *in silico* data analysis. In the last decade, many breakthroughs focused on sample preparation, leading to more efficient protein extraction protocols such as Filter Aided Sample Preparation (FASP) (Wiśniewski, Zougman, et al. 2009), Single-Pot Solid-Phase-Enhanced Sample Preparation (SP3) (Hughes et al. 2014), Suspension Trapping (STrap) (Zougman et al. 2014) and detergent-free method known as Sample Preparation by Easy Extraction and Digestion (SPEED) (Doellinger et al. 2020). Meanwhile, mass-spectrometry and data management constantly improved, supported by development of the Orbitrap technology (Hu et al. 2005) and impressive growth of computational power, allowing deeper and quicker acquisition of proteomic data. However, complex samples are still challenging to tackle. Culture-independent bacterial identification by proteo-typing in complex matrices such as blood or urine using tandem mass-spectrometry for clinical diagnosis is in its infancy (Roux-Dalvai et al. 2019; Kondori et al. 2021), and very few dual-proteomics research studies investigating complex host-pathogen interactions have been published (Geddes-McAlister et al. 2021; Ball et al. 2020; Willems et al. 2021; Leseigneur et al. 2022; Masson et al. 2021). Additionally, research on highly virulent microorganisms is restricted by regulatory requirements for safety purposes. Mandatory sample treatment steps generally include pathogen inactivation and pathogen DNA and RNA elimination in biosafety level (BSL) −3 or −4 laboratories.

*Yersinia pestis* is the highly virulent bacillus responsible for plague, a quickly fatal human disease if left untreated. This reemerging disease whose epidemics had a considerable death toll in human history generally manifests as bubonic plague following bites from fleas infected with *Y. pestis*. Primary pulmonary plague is the second form of the disease, which occurs by inhalation of infected respiratory droplets or aerosols usually emitted by an infected patient. In some cases, injection of *Y. pestis* by fleas directly into the bloodstream leads to so-called septicemic plague (Perry et al. 1997). In the three forms of the disease, bacteria reach the blood resulting in systemic infection. While plague pathogenesis has been investigated for decades, the bacteremic phase of the disease remains to be fully deciphered.

Here, we report the design of experimental pipelines allowing characterization of human and bacterial proteomes in plasma and blood infected with *Y. pestis*. We first applied protein extraction methods to *Y. pestis* comparing the SPEED method (Doellinger et al. 2020) based on trifluoracetic acid (TFA) lysis, the standard bead-beating lysis in urea followed by the FASP method (Wiśniewski, Zougman, et al. 2009), and sonication in sodium dodecyl sulfate (SDS) followed by buffer exchange and the FASP method (Wiśniewski, Zougman, et al. 2009). SPEED surpassed the other methods, allowing quick, efficient and safe sample preparation in BSL-3 environment with fully virulent *Y. pestis*. It was thus further used to conduct the first dual proteomic study of a bacterial species in human plasma and human blood, revealing the pathogenic proteomic signature of *Y. pestis*.

## Results

### Comparison of urea-, SDS- and TFA-based methods for *Y. pestis* proteomics

To study the *Y. pestis* proteome, we first used the avirulent *Y. pestis* CO92 strain lacking pCD1 in a BSL-2 environment. Bacteria were grown on lysogeny broth with 0.002% pork hemin (LBH) agar plates at 37°C for 24 hours. We compared the three following protein extraction protocols: i) urea-based method: bacterial lysis by bead beating in 8M urea buffer, followed by reduction-alkylation and digestion using the FASP method, ii) SDS-based method: bacterial resuspension in 4% SDS buffer, heating at 95°C and lysis by sonication, followed by buffer exchange with 8M urea, reduction-alkylation and digestion using the FASP method, and iii) TFA-based SPEED method: bacterial lysis upon TFA acidification, followed by neutralization with 10 volume of 2M Tris base, reduction-alkylation and digestion.

Three protein quantification methods were compared to assess their compatibility with urea, SDS or TFA-Tris buffers. Bradford (Bradford 1976), BCA (Smith et al. 1985) and Qubit assays were only compatible with one buffer each, and not with 10% TFA-2M Tris base buffer (Table 1). Consequently, we implemented protein quantification by tryptophan fluorescence spectroscopy (Wiśniewski and Gaugaz 2015) which was compatible with the three buffers (Table 1, Fig EV1A).

**Table 1.** Compatibility of Bradford, BCA, Qubit and tryptophan fluorescent assay with 8M Urea, 4% SDS and 10% TFA - 2M Tris lysis buffer for protein quantification.

|  |  | Extraction method |  |  |
| --- | --- | --- | --- | --- |
|  |  | 8M Urea | 4% SDS | 10% TFA - 2M Tris |
| **Measure**<br>**method** | **Bradford** | Compatible | Incompatible | Incompatible |
|  | **BCA** | Incompatible | Compatible | Incompatible |
|  | **Qubit** | Compatible | Incompatible | Incompatible |
|  | **Tryptophan fluorescence** | Compatible | Compatible | Compatible |

After data-dependent acquisition (DDA) analysis of triplicates using an Orbitrap massspectrometer, we could detect a total of 1,587 proteins with at least one of the protocols, 1,390 bacterial proteins with the urea-based protocol, 1,444 proteins with the SDS-based protocol and 1398 proteins with the TFA-based protocol (Fig EV1B, Table EV1). Overall 1,194 proteins were common to the three protocols, 30 proteins were only detected using the urea-based protocol, 72 proteins were specifically detected using the SDS-based protocol and 34 proteins were exclusive to the TFA-based protocol (Fig EV1B). A total of 189 proteins were detected using urea- or SDS-based protocols but not by using the TFA-based protocol. While the three protocols led to similar *Y. pestis* proteomes, the TFA-based protocol surpassed both bead-beating and sonication methods, which required a fastidious centrifugation step, by its quick implementation, high-throughput potential and application to very small amount of biomass, which is important when dealing with precious purified fractions from complex samples. Moreover, the minimal number of steps of the SPEED method reduced the risk of experimental errors and variability. These benefits led us to choose the SPEED method to further investigate *Y. pestis* proteome in a BSL-3 environment.

### Inactivation of virulent *Y. pestis* using the SPEED method in BSL-3 environment

Mass spectrometric analysis of virulent *Y. pestis* proteome performed in BSL-1 platform requires bacterial inactivation according to French regulatory protocols for “Micro-Organismes et Toxines” (MOT) from the Agence Nationale de Sécurité du Médicament et des Produits de Santé (ANSM). *Y. pestis* CO92 was grown on LBH agar plates at 37°C for 24 hours. Bacterial proteins were extracted using the SPEED method. To validate absence of viable bacteria, lysogeny broth (LB) was inoculated with TFA lysates before and after neutralization and incubated under agitation at 28°C for 72 hours. Absence of growth measured by optical density at 600 nm certified inactivation of viable bacteria in both samples. To assess absence of bacteriostatic effect of the inactivating agent, the LB samples were spiked with 1 to 2 bacteria and incubated 72 hours at 28°C. Growth measured at 600 nm revealed absence of bacteriostatic effect of the inactivating agent. *Y. pestis* specific phage assay was used to assess the absence of contamination of the spiked sample. To fulfill the MOT regulation, bacterial DNA degradation was assessed by PCR on *Y. pestis caf1*, *yopM*, and *pla* genes after a 30-minutes dialysis to remove inhibitory salts. While amplicons corresponding to the three genes were detected when PCR was performed on *Y. pestis* DNA used as positive control, they could not be detected when PCR was performed on SPEED samples. Together, our results indicate that the SPEED method allows bacterial inactivation and DNA degradation directly after bacterial lysis and, consequently, that the following reduction, alkylation and digestion steps and mass-spectrometry can be performed in a BSL-1 environment.

### Proteomic reshaping of *Y. pestis* grown in human plasma

To identify *Y. pestis* proteins expressed in conditions mimicking the human bacteremic phase of plague, we first compared the proteomes of bacteria grown in human plasma or in culture media. Proteomes of the fully virulent *Y. pestis* CO92 strain were characterized in six BSL-3 culture conditions (Fig 1A): i) inoculum grown overnight on LBH agar at 25°C (I25), ii) inoculum grown overnight on LBH agar at 37°C (I37), iii) growth in chemically defined medium M9 at 37°C inoculated with I25 (I25-M9), iv) growth in M9 at 37°C inoculated with I37 (I37-M9), v) growth in human plasma at 37°C inoculated with I25 (I25-Plasma) and vi) growth in human plasma at 37°C inoculated with I37 (I37-Plasma). The two different inoculum temperatures reflect the environmental origin of bacteria entering the bloodstream: 25°C mimics direct transmission from fleas in primary septicemic plague and 37°C mirrors hematogenous dissemination from the bubo or lungs in secondary septicemic plague.

**Figure 1.**
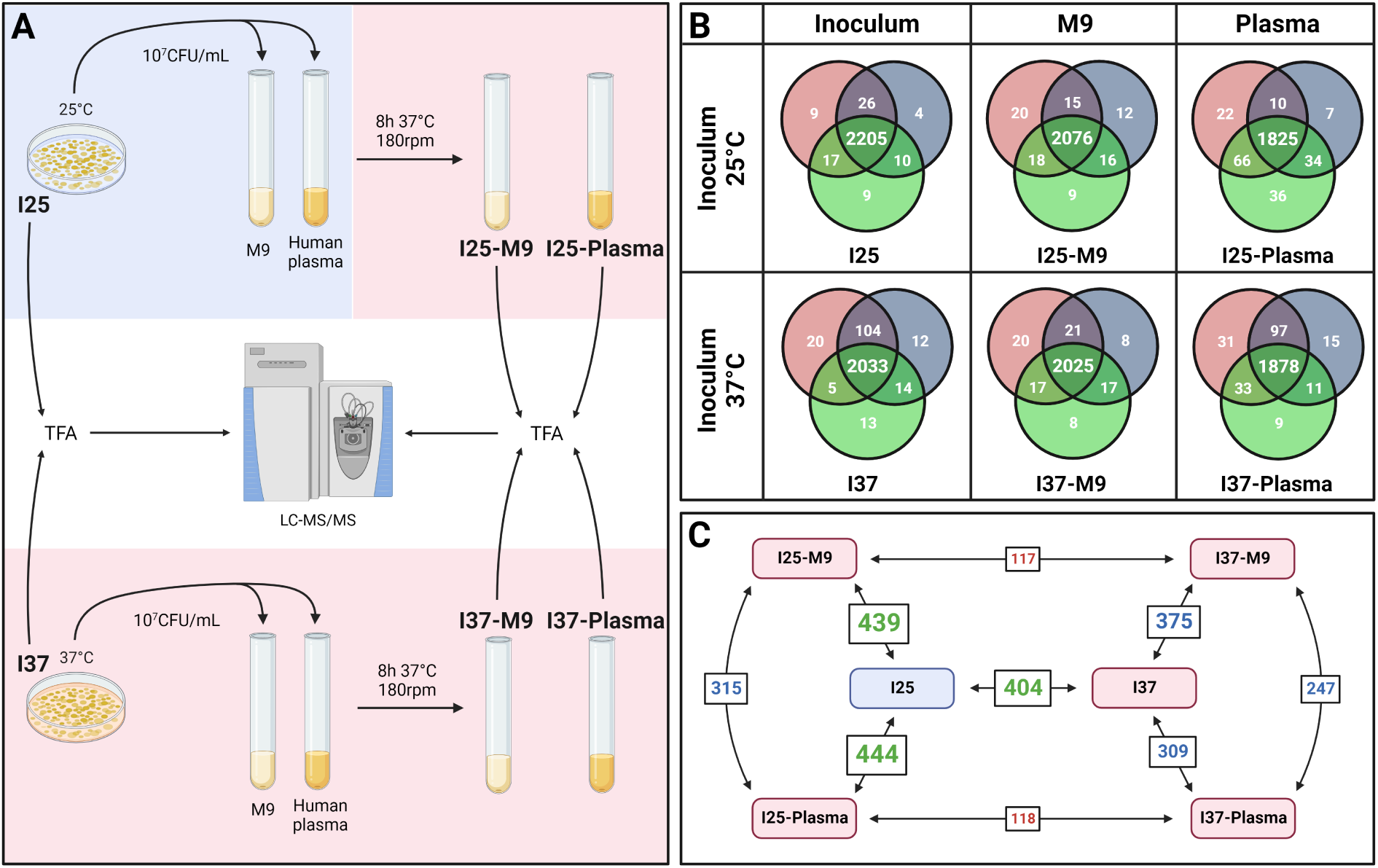
(A) Protocol comparing *Y. pestis* grown on LBH plate at 2 different temperatures and after growth in M9 or human plasma at 37°C. (B) Venn diagrams of the number of identified proteins across replicates in the same condition, for samples of *Y. pestis* grown on LBH, in M9 or human plasma at different temperatures. Each circle in the Venn diagrams corresponds to a replicate. (C) Number of differentially abundant proteins between samples after *Y. pestis* growth on LBH, in M9 or human plasma at different temperatures. These numbers only consider the proteins detected in both conditions. Condition in the blue rectangle is grown at 25°C and conditions in the red rectangles are grown at 37°C. Numbers in green are associated with change in temperature culture, numbers in blues are associated with a change in media grown at 37°C, and numbers in red are associated with a change in inoculum temperature in the same medium.

We identified 2,382 bacterial proteins in at least one sample across all conditions, from a total of 3,915 coding sequences and small open reading frame-encoded proteins (Table EV2) (Cao et al. 2021; Parkhill et al. 2001). Optimization of Orbitrap-based mass-spectrometry in DDA mode led to a gain in the number of proteins detected compared to our initial proteomic analysis comparing the extraction protocols, which identified only 1,587 proteins (see above). Between 2,025 to 2,205 proteins were identified in LBH and M9 samples (Fig 1B). The number of protein identifications was reduced to 1,825-1,878 in plasma samples, presumably due to presence of human proteins (Fig 1B). The most extensive detection was achieved in the LBH sample grown at 25°C in which 2,205 proteins were identified in the three replicates compared to the 2,033 proteins detected in LBH samples grown at 37°C. Detection of proteins of lower abundance could have been prevented at 37°C by the presence of highly produced proteins in the samples at that temperature, such as the F1 pseudocapsule (Table 2) and components of the type three secretion system (T3SS) (Demeure et al. 2019) (Table 3).

**Table 2.**
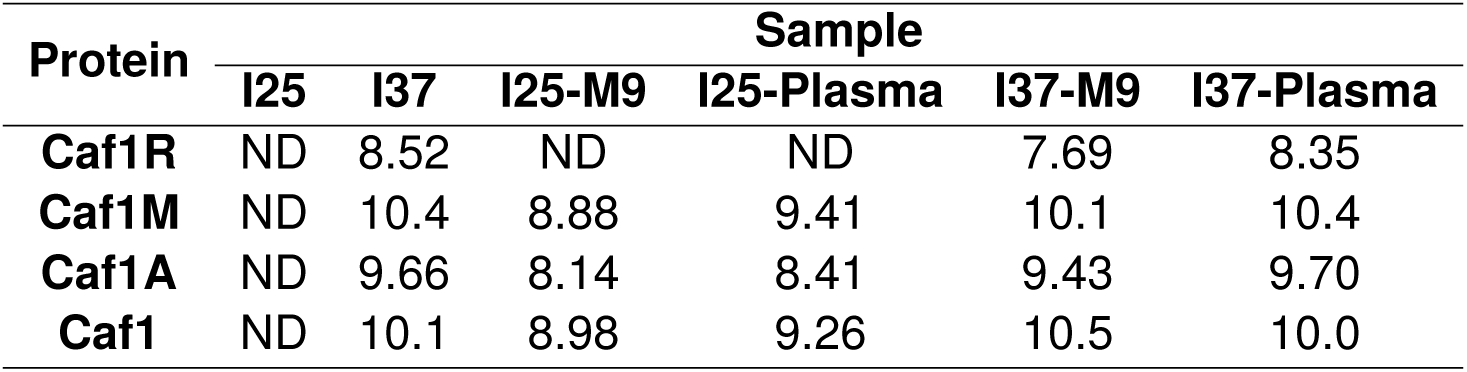
Value of log_10_(LFQ) for the *caf* operon encoding *Y. pestis* pseudocapsule. ND: Not Detected.

| Protein | Sample |  |  |  |  |  |
| --- | --- | --- | --- | --- | --- | --- |
|  | I25 | I37 | I25-M9 | I25-Plasma | I37-M9 | I37-Plasma |
| **Caf1R** | ND | 8.52 | ND | ND | 7.69 | 8.35 |
| **Caf1M** | ND | 10.4 | 8.88 | 9.41 | 10.1 | 10.4 |
| **Caf1A** | ND | 9.66 | 8.14 | 8.41 | 9.43 | 9.70 |
| **Caf1** | ND | 10.1 | 8.98 | 9.26 | 10.5 | 10.0 |

**Table 3.** Log_2_(fold change) of protein abundance for proteins involved in the type 3 secretion system (T3SS). A value of “15” is indicated for proteins only detected in the first condition and a value of “-15” is indicated for proteins only detected in the second condition.

| Locus | Protein | Comparison |  |  |  |  |
| --- | --- | --- | --- | --- | --- | --- |
|  |  | I25 | I37 | I37 | I25-M9 | I37-M9 |
|  |  | vs<br>I37 | vs<br>I37-M9 | vs<br>I37-Plasma | vs<br>I25-Plasma | vs<br>I37-Plasma |
| YPCD1.05c | SycE | -6.6 | -1.1 | 0.1 | 1.7 | 1.2 |
| YPCD1.06 | YopE | -7.7 | -2.0 | -0.1 | 2.8 | 1.9 |
| YPCD1.19c | YopK | -15.0 | -2.2 | 0.7 | 4.3 | 2.9 |
| YPCD1.20 | YopT | -15.0 | -2.2 | 15.0 | 0.0 | 15.0 |
| YPCD1.21 | SycT | 0.0 | -15.0 | 0.0 | 0.0 | 15.0 |
| YPCD1.26c | YopM | -15.0 | -0.8 | 1.2 | 2.0 | 2.0 |
| YPCD1.28c | YopD | -9.3 | -1.3 | 0.8 | 2.5 | 2.2 |
| YPCD1.29c | YopB | -15.0 | -1.6 | 1.0 | 3.9 | 2.7 |
| YPCD1.30c | LcrH | -15.0 | -1.4 | 0.5 | 2.8 | 1.9 |
| YPCD1.31c | LcrV | -8.5 | -1.3 | 0.9 | 3.6 | 2.2 |
| YPCD1.32c | LcrG | -9.2 | -1.5 | 1.6 | 4.8 | 3.1 |
| YPCD1.33c | LcrR | -15.0 | -2.0 | 15.0 | 15.0 | 15.0 |
| YPCD1.34c | LcrD | -15.0 | -2.0 | 0.9 | 3.0 | 2.9 |
| YPCD1.35c | YscY | -15.0 | -1.7 | 1.4 | 3.0 | 3.1 |
| YPCD1.36c | YscX | -15.0 | -2.0 | 0.8 | 3.1 | 2.9 |
| YPCD1.37c | SycN | -6.5 | -2.5 | 0.0 | 2.7 | 2.5 |
| YPCD1.38c | TyeA | -15.0 | -1.6 | 0.3 | 2.7 | 1.9 |
| YPCD1.39c | YopN | -15.0 | -0.1 | 0.8 | 0.9 | 0.9 |
| YPCD1.40 | YscN | -15.0 | -1.7 | 1.0 | 3.5 | 2.7 |
| YPCD1.41 | YscO | -15.0 | -2.5 | 1.8 | 4.5 | 4.3 |
| YPCD1.42 | YscP | -15.0 | -1.5 | 1.4 | 4.4 | 2.9 |
| YPCD1.43 | YscQ | -15.0 | -0.9 | 1.3 | 2.7 | 2.2 |
| YPCD1.44 | YscR | 0.0 | -15.0 | 0.0 | 15.0 | 15.0 |
| YPCD1.46 | YscT | 0.0 | -15.0 | 0.0 | 15.0 | 15.0 |
| YPCD1.47 | YscU | -15.0 | -2.2 | 1.0 | 3.2 | 3.2 |
| YPCD1.48 | YscW | -15.0 | -0.8 | -0.3 | 1.2 | 0.5 |
| YPCD1.49 | VirF | -15.0 | -1.1 | 0.2 | 2.7 | 1.3 |
| YPCD1.51 | YscB | -9.2 | -1.7 | 0.7 | 2.7 | 2.4 |
| YPCD1.52 | YscC | -15.0 | -2.6 | 0.7 | 5.0 | 3.4 |
| YPCD1.53 | YscD | -15.0 | -2.8 | 0.9 | 5.7 | 3.6 |
| YPCD1.54 | YscE | -4.0 | -2.2 | -0.2 | 2.6 | 2.0 |
| YPCD1.55 | YscF | -6.2 | -2.5 | 0.3 | 3.7 | 2.8 |
| YPCD1.56 | YscG | -3.9 | -2.3 | -0.1 | 3.2 | 2.2 |
| YPCD1.57 | YopR | -15.0 | -2.6 | 0.7 | 15.0 | 3.4 |
| YPCD1.58 | LcrO | -15.0 | -3.7 | 1.6 | 15.0 | 5.1 |
| YPCD1.59 | YscJ | -15.0 | -3.4 | 0.3 | 4.5 | 3.6 |
| YPCD1.60 | YscK | -6.4 | -1.7 | 0.8 | 3.5 | 2.5 |
| YPCD1.61 | YscL | -15.0 | -1.6 | 0.5 | 3.0 | 2.1 |
| YPCD1.62 | LcrQ | -3.5 | -0.5 | 0.0 | -1.4 | 0.5 |
| YPCD1.67c | YopH | -5.2 | -1.8 | 1.4 | 3.6 | 3.2 |
| YPCD1.71c | YopJ | -15.0 | -1.2 | 1.4 | 6.3 | 2.6 |
| YPCD1.72c | YopO | -15.0 | -0.6 | 1.0 | 2.1 | 1.5 |
| YPCD1.95c | SycH | -1.5 | 1.0 | 0.5 | -0.6 | -0.5 |

Comparison of differential abundances of the proteins between samples revealed extensive proteome reshaping across conditions (absolute fold change *>* 2 and adjusted p-value *<* 0.01, Fig 1C). Expectedly, a shift in temperature and culture medium (I25 vs I25-M9, I25 vs I25-Plasma) was associated with the highest number of differentially abundant proteins, with 439 and 444 proteins (taking into account only the proteins detected in both compared conditions), respectively (Fig 1C). Temperature shift alone also induced major proteome changes when I25 was compared to I37 (404 differentially abundant proteins). A change of culture medium (I37 vs I37-M9, I37 vs I37-Plasma, I37-M9 vs I37-Plasma) led to slightly fewer differentially abundant proteins with 375, 309, 247 proteins, respectively (Fig 1C). More modest changes were observed when only shifting the inoculum temperature in M9 (117 proteins for I25-M9 vs I37-M9) and in plasma (118 proteins for I25-Plasma vs I37-Plasma). A full description of numbers of differentially abundant proteins between each condition, as well as numbers of proteins present in one condition but absent in another are indicated in Fig EV2. To assess the influence of donors on the proteome of *Y. pestis* grown in plasma, we compared proteomes of bacteria grown in the plasma of four different donors inoculated with I25 and I37 in triplicates. For each donor, we identified between 1,486 and 1,696 proteins common to the three replicates (Fig EV3, Table EV3), slightly different numbers than in our initial proteome analysis (see above, Fig 1B). Comparison of samples derived from the four different donors identified approximately 70 proteins (37°C inoculum) to 100 proteins (25°C inoculum) which were differentially abundant, showing a good reproducibility of proteomes when *Y. pestis* was grown in plasma of different donors.

Together, our results reveal a major *Y. pestis* protein expression remodeling upon growth in human plasma at 37°C.

### Dual proteomics of *Y. pestis* in human blood

To refine our model of bacteremia *ex vivo*, we investigated *Y. pestis* grown in whole human blood. A previous proteotyping study used blood cell lysis to deepen detection of bacterial peptides and identify *Staphylococcus aureus*, *Escherichia coli* and *Candida albicans* for clinical diagnosis (Kondori et al. 2021). Although this protocol enables proteomic analysis of bacteria in whole blood, it precludes identification of the human proteome, which is useful to understand the host response to infection. Thus, we set up an original pipeline allowing to analyze both bacterial and human proteomes in whole blood. We first compared growth of *Y. pestis* in whole human blood using ethylenediaminetetraacetic acid (EDTA), acid-citrate-dextrose (ACD) or heparin as anticoagulant. We selected ACD, as *Y. pestis* grew the best in this blood compared to heparin- and EDTA-anticoagulated blood, reaching 10^8^ bacteria/mL after an 8 hour-incubation (Fig EV4). Human blood is a highly complex biological fluid characterized by a high concentration of red blood cells (*≈* 4.5-12.10^9^ cells/mL), and lower concentrations of platelets (*≈* 150-475.10^6^ cells/mL) and white blood cells (*≈* 5-10.10^6^ cells/mL). Neutrophils constitute the vast majority of white blood cells (40-60%), followed by lymphocytes (20-40%) and monocytes (2-8%), eosinophils ranging from 1 to 4% and basophils 0.5 to 1%. The high number of human cells compared to bacteria in our sample (10^8^ bacteria/mL) and the vast dynamic range of human proteins abundance (*≈* 10^13^), make characterization of dual proteomes very challenging. We thus set up a pipeline to enrich *Y. pestis* associated to relevant immune cells, neutrophils and monocytes (Arifuzzaman et al. 2018; Pujol et al. 2005; Spinner et al. 2014; Dudte et al. 2017; Osei-Owusu et al. 2019). After 8-hour incubation of bacteria in ACD-anticoagulated human blood, we purified these cells by immunomagnetic separation (Fig 2). Using CD66b neutrophil-specific antibodies and CD14 monocyte-specific antibodies, 2.10^5^ viable neutrophils and 6.10^4^ viable monocytes were recovered per milliliter of blood, which account for around 10% of the total neutrophil and monocyte populations. Besides technical issues, cell loss could be due to reduced lifespan of neutrophils *ex vivo* and *Y. pestis* capacity to induce cell death (Marketon et al. 2005; Osei-Owusu et al. 2019). From 10^8^ bacteria per milliliter of whole blood, 4.10^6^ bacteria were associated to the neutrophils fraction (1/25^th^ of total bacteria), and 5.10^5^ bacteria were associated to the monocyte fraction (1/200^th^ of total bacteria). This approach allowed to lower sample complexity and to switch from the initial ratio of 10^8^ bacteria to 5.10^9^ human cells (1:50), to ratios of 4.10^6^ bacteria to 2.10^5^ neutrophils (20:1) and 5.10^5^ bacteria to 6.10^4^ monocytes (8:1). We then extracted proteins using the SPEED method and analyzed samples by mass-spectrometry using a DIA mode (Fig 2). To this end, we created two spectral libraries. The first library was constructed using the antibody-mediated cell enrichments combined with a Nanotrap^®^ enrichment, which had previously been shown to concentrate *Y. pestis* bacteria from whole blood and bacterial proteins from lysates (Ii et al. 2021). The second library was constructed from *Y. pestis* grown in LB broth at 37°C to obtain the deepest peptide library. We identified 1,366 bacterial and 4,187 human proteins in the neutrophil fraction, and 877 bacterial and 3,969 human proteins in the monocyte fraction (Table EV4). It is, to our knowledge, the first report of a pipeline allowing semi-quantitative dual proteomics extracted from a complex matrix such as infected whole blood, which could moreover be applied in a BSL-3 environment.

**Figure 2.**
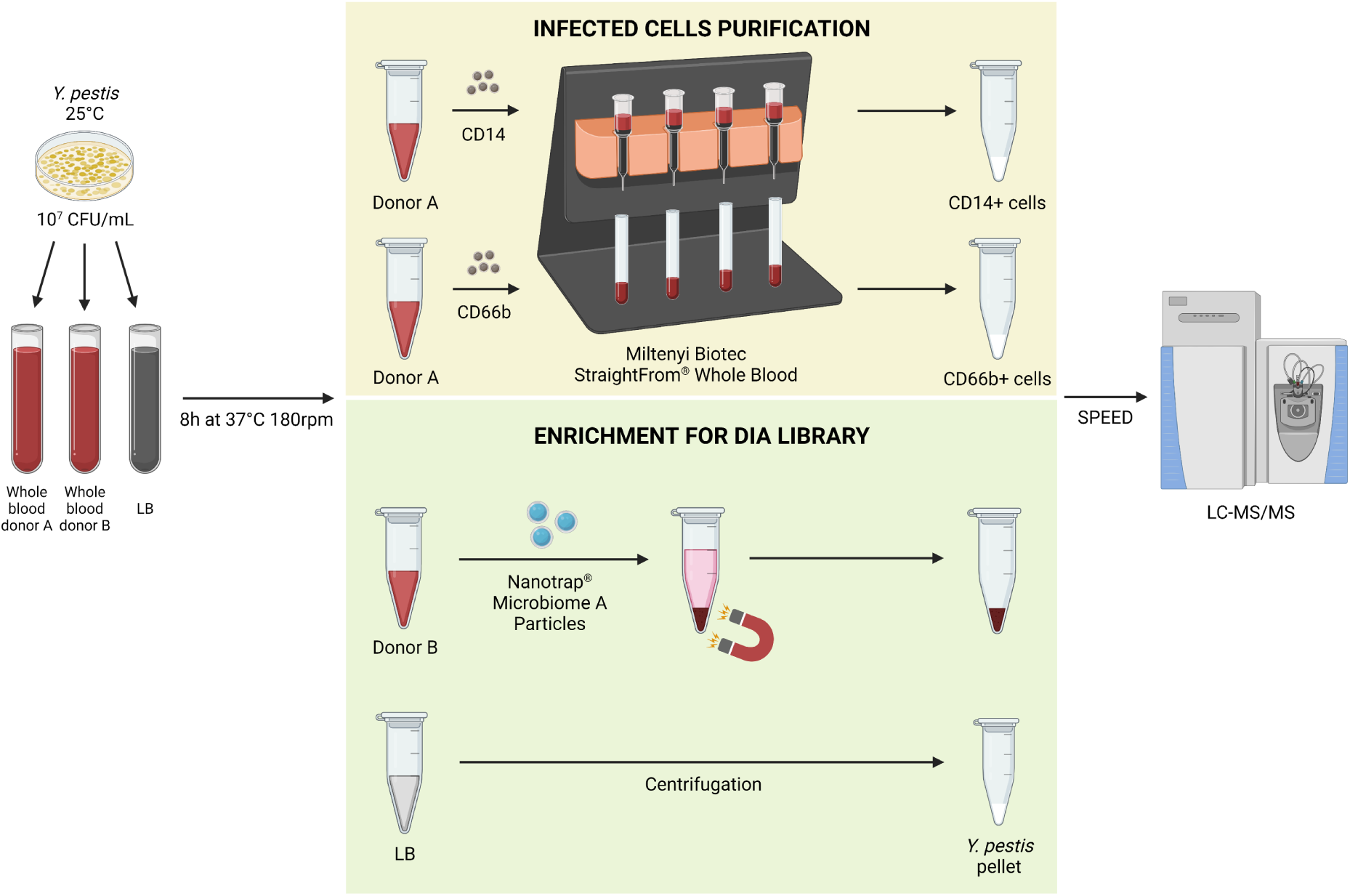
Workflow for enrichment of cell-associated *Y. pestis* from human whole blood, and library construction based on Nanotrap enrichment and pure culture in LB.

### Library depths and normalization strategies

Nanotrap enrichment led to a good peptide coverage of bacterial proteins that were very abundant in blood, such as MetE, MetF or MetR, which were also enriched when *Y. pestis* was grown in human plasma (Table 4). Similarly, T3SS proteins, such as YscX (Gurung et al. 2022; Day et al. 2000) and YopK, YopB, YopD, YopT, YopH, whose secretion is induced in contact to host cells (Osei-Owusu et al. 2019) were identified. We could identify more than 2,800 bacterial proteins during the secondary library construction consisting in a DDA analysis of the fractionated pool of *Y. pestis* grown in LB. To our knowledge, it is the most complete proteome of a *Yersinia* species. DIA analysis led to identification of 1,366 bacterial proteins in the neutrophil fraction, 877 bacterial proteins in the monocyte fraction, 1,681 bacterial proteins in the Nanotrap enrichment samples, and 2,335 bacterial proteins in *Y. pestis* grown in LB (Table 5). Hierarchical clus-tering (Fig 3A) and principal component analysis (Fig 3B) based on MS peaks associated to bacterial and human proteins showed an excellent clustering of the replicates as well as an enrichment of bacterial and human proteins upon Nanotrap capture. The smaller number of bacterial proteins identified in the monocyte fraction compared to the neutrophil fraction (Table 5 and Fig 3A) are in line with the number of bacteria per cell, ranging from 8 bacteria per cell in monocyte to 20 bacteria per cells for the neutrophils (see above).

**Figure 3.**
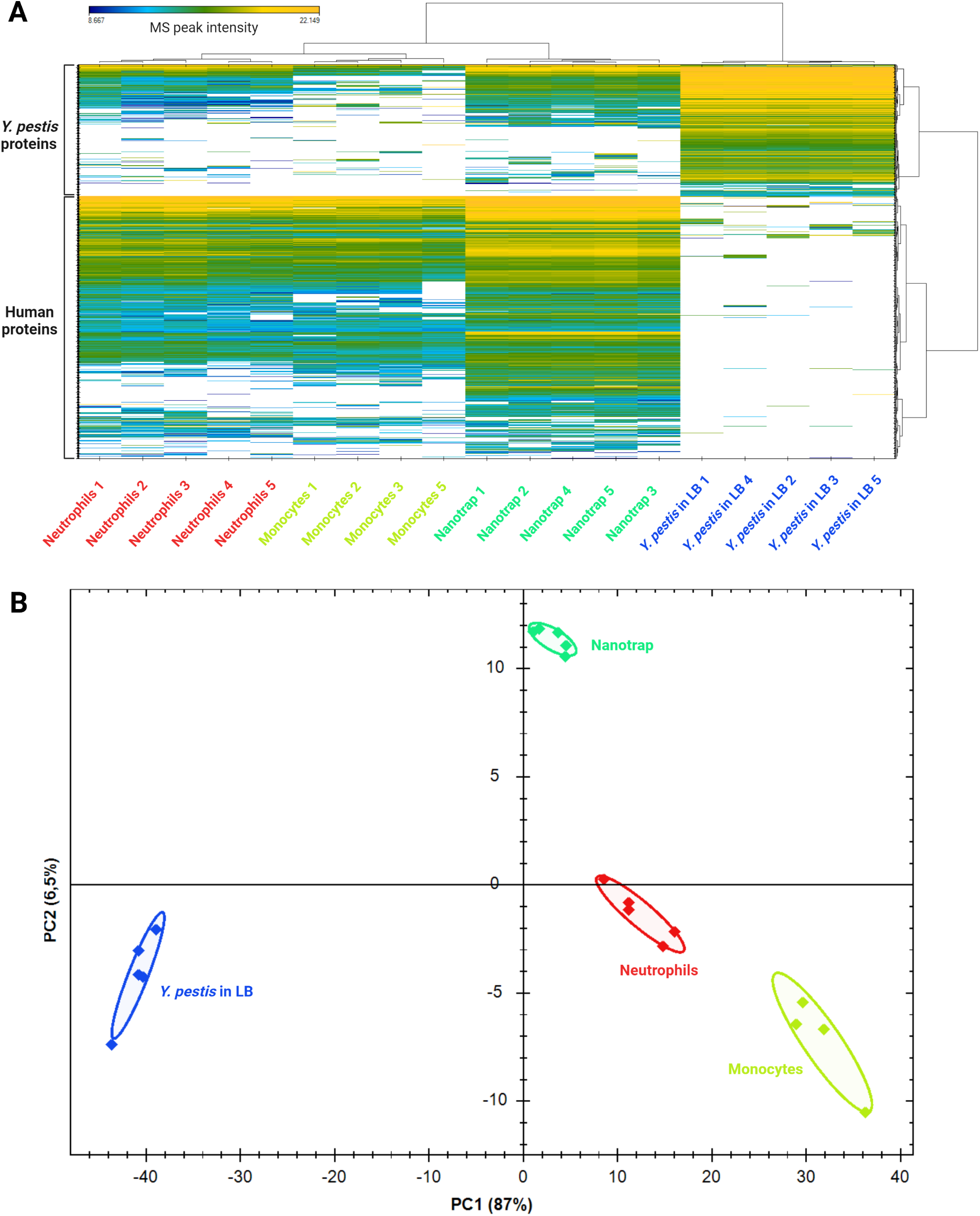
(A) Heatmap and (B) principal component analysis (PCA) based of the MS peak intensities for the bacterial and the human proteins on the neutrophil fraction, the monocyte fraction, the Nanotrap enrichment and the *Y. pestis* pure culture in LB. The replicate 4 of the monocyte fraction was removed due to a contamination with red blood cells during cell purification.

**Table 4.**
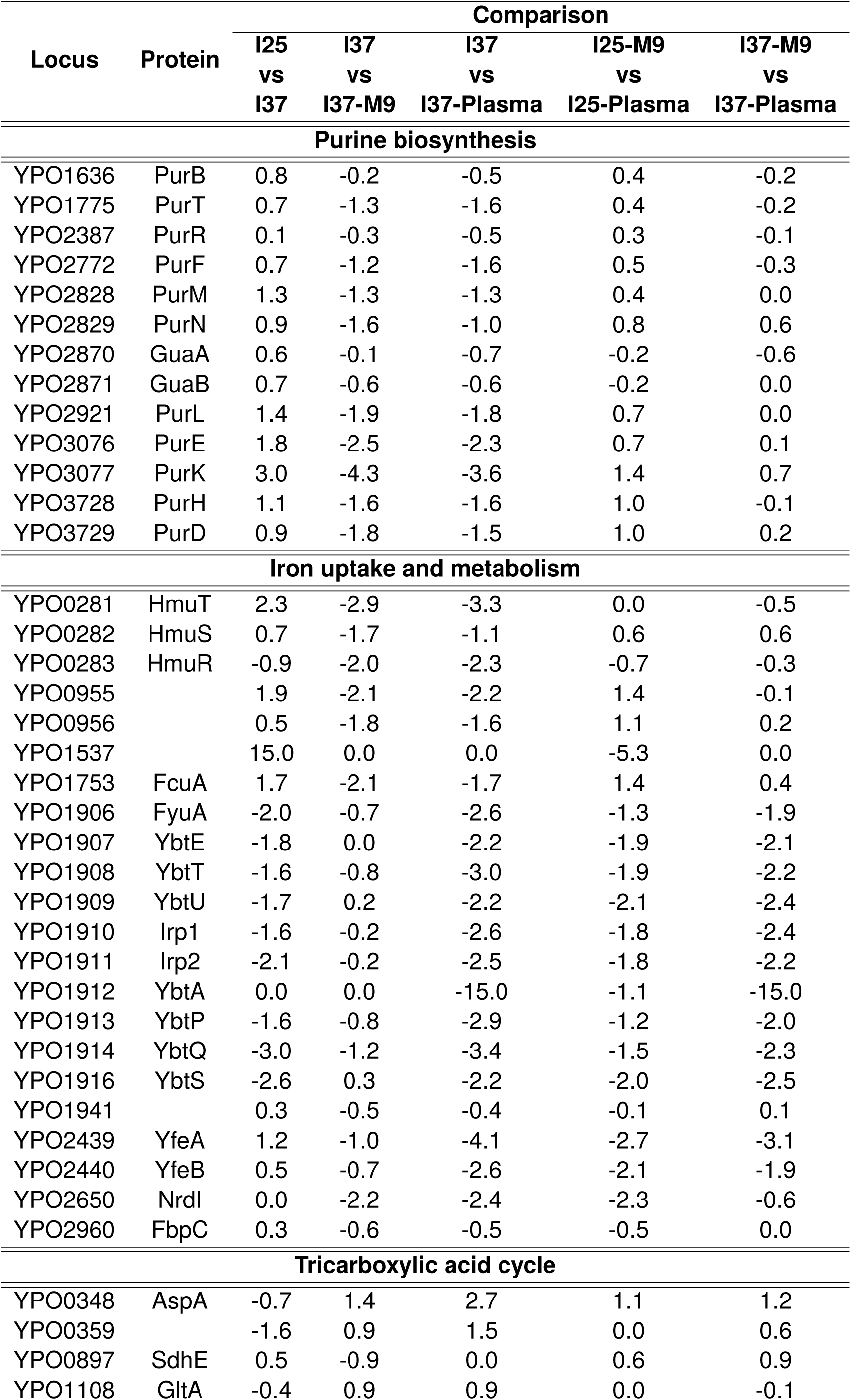
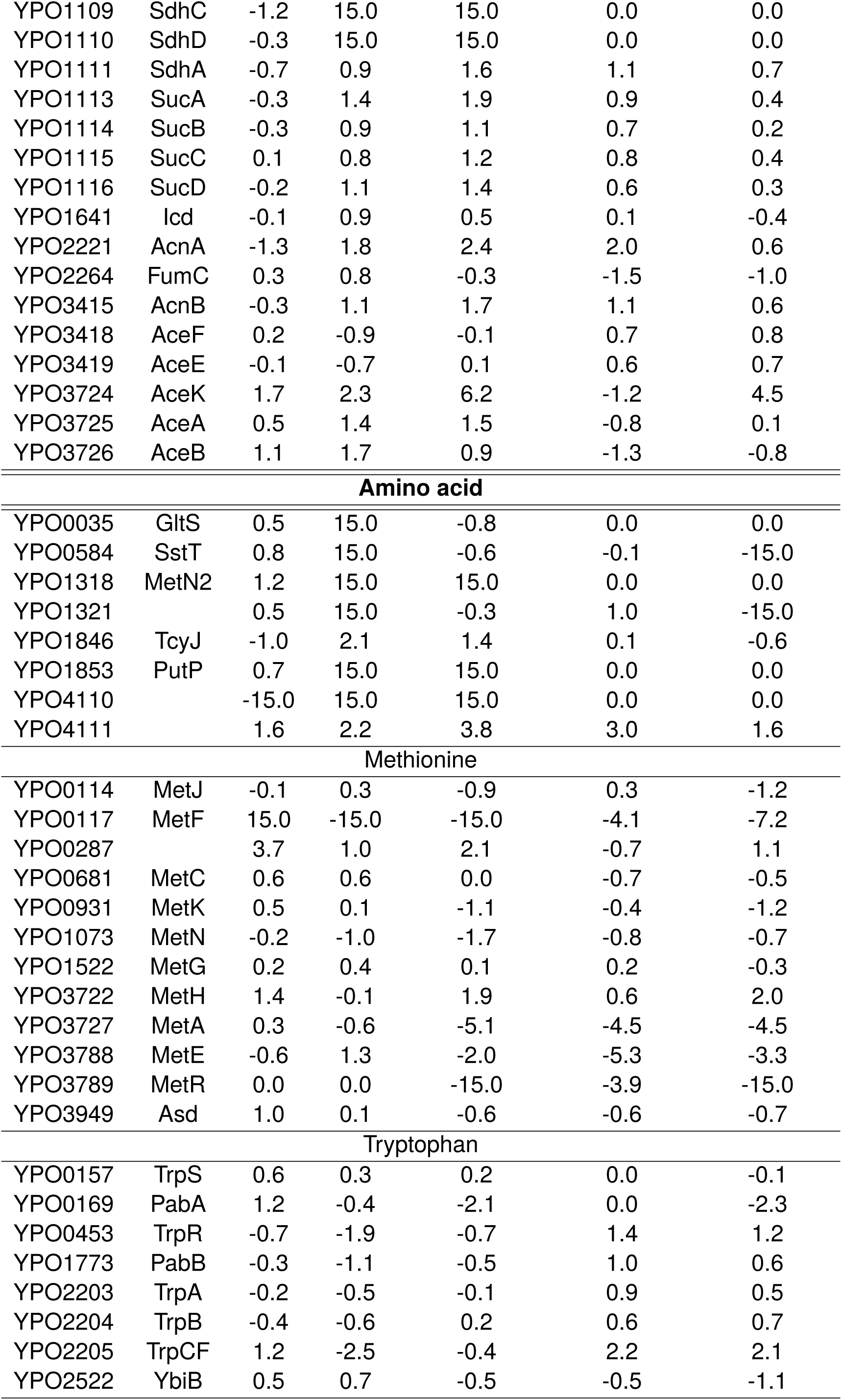
Log_2_(fold change) of protein abundance for proteins involved in metabolism and iron uptake. A value of “15” is indicated for proteins only detected in the first condition and a value of “-15” is indicated for proteins only detected in the second condition.

**Table 5.**
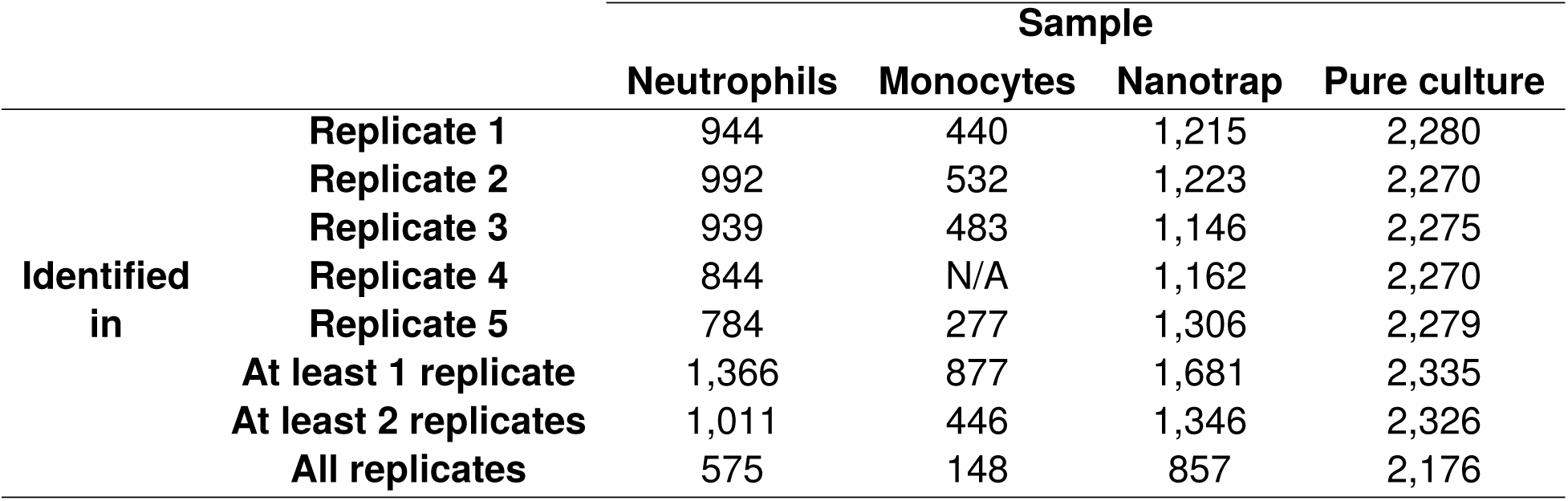
Number of bacterial proteins identified after the DIA analysis of the blood fractions, the Nanotrap enrichment and the pure culture in LB. The replicate 4 of the monocyte fraction was removed due to a contamination with red blood cells during cell purification.

We tested several normalization strategies to take into account the different ratios of bacteria per cell in both fractions. These strategies are detailed in Table 6. The differential analysis was performed taking into account either the proteins identified in at least one sample, strategies A (all), H (human), Ya (*Yersinia* all) and Yc (*Yersinia* per condition), which allows to identify proteins which are detected in one condition but not the other, or in at least two samples per condition, strategies A2 (all), H2 (human), Ya2 (*Yersinia* all) and Yc2 (*Yersinia* per condition), which reduces the number of protein considered in the analysis but also reduces the rate of false positive identification. Most of the differential analysis below will thus result from this second set of strategies.

**Table 6.** Normalization and differential analysis strategies for whole blood fraction analysis. The number of samples where peptides were found (last column) was only taken into account during the differential analysis and not during normalization, which was based on all peptides. Yp: *Y. pestis*. Hsa: *Homo sapiens*. Spl: sample. Cond: biological condition (monocyte or neutrophil fraction).

| Name | Description | Dataset | Median normalization | Normalized on | Prot. quantified in at least 2 spl |
| --- | --- | --- | --- | --- | --- |
| A | All | Hsa+Yp | on all spl | Hsa | in a cond |
| A2 | All (2 spl) | Hsa+Yp | on all spl | Hsa | in each cond |
| H | Hsa | Hsa | on all spl | Hsa | in a cond |
| H2 | Hsa (2 spl) | Hsa | on all spl | Hsa | in each cond |
| Ya | Yp all | Yp | on all spl | Yp | in a cond |
| Ya2 | Yp all (2 spl) | Yp | on all spl | Yp | in each cond |
| Yc | Yp by cond | Yp | by cond | Yp | in a cond |
| Yc2 | Yp by cond (2 spl) | Yp | by cond | Yp | in each cond |

The A2 strategy consisted in normalizing the intensities of both human and bacterial proteins by equalizing the medians of the human proteins in the samples across the two conditions, and to perform a differential analysis on proteins which were detected in at least 2 replicates per condition. When using this strategy, the lower ratio of bacteria per cells in the monocyte fraction compared to the neutrophil fraction was in line with the analysis of the differential abundance of the bacterial proteins. Indeed, a GSEA on biological processes GO terms and KEGG pathways revealed an enrichment of translation and ribosomal processes in the neutrophil fraction, considered here as house-keeping processes, and the associated enriched protein log_2_(fold change) were centered around 2.5 (5.6 fold change) (Fig 4A and B). This effect was also observed when we normalized only the bacterial proteins based on the median of each conditions separately (strategy Yc2, Fig 4C), resulting in a volcano plot where a large set of proteins mainly corresponding to ribosomal proteins were uniformly upregulated in the neutrophil fraction (Fig 4C). The calculated 5.6-fold ratio based on ribosomal proteins was higher than the previous calculated ratio based on bacterial enumeration (8 viable bacteria per monocyte and 20 viable bacteria per neutrophil, resulting in a 2.5-fold ratio) which could be explained by the lower intracellular survival of *Y. pestis* in neutrophils compared to monocytes (Arifuzzaman et al. 2018; Pujol et al. 2005; Spinner et al. 2014; Dudte et al. 2017; Osei-Owusu et al. 2019). When normalizing only the bacterial proteins by equalizing the median of the samples from the two conditions taken together (strategy Ya2), we observed a shift of the ribosomal proteins to the non differentially abundant group when performing an enrichment analysis on non differentially abundant proteins (Table 7). As the number of identified proteins in the monocyte was much lower than in neutrophils, due to the lower number of bacteria per cell and subsequent reduced sensitivity, most of the less abundant proteins in the monocyte fraction could presumably not be detected. Only upregulated proteins in the monocyte fraction, identified in both the monocyte and neutrophil fractions, could be measured and compared, resulting in an asymmetrical volcano plot (Fig 4D).

**Figure 4.**
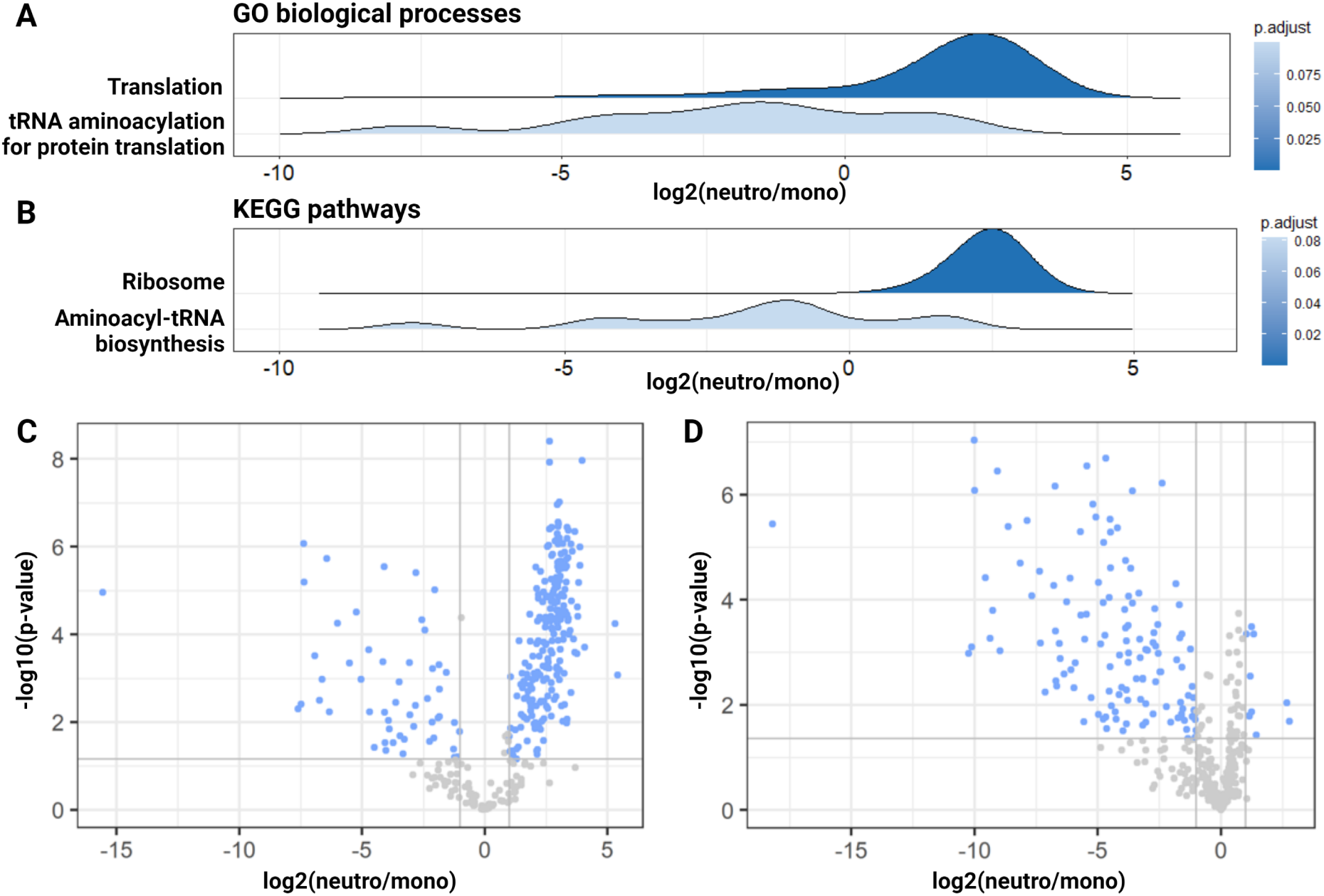
(A-B) Ridgeplots of enriched (A) GO biological processes or (B) KEGG pathways after performing a GSEA on *Y. pestis* protein differential abundances between the neutrophil and monocyte fractions using the A2 normalization strategy. The curve represents the density of proteins in the enriched terms depending on the fold change of these proteins. (C-D) Volcano plots showing differentially abundant *Y. pestis* proteins between the neutrophil fraction and the monocyte fraction using (C) the Yc2 strategy and (D) the Ya2 strategy. In blue are the proteins which are significantly differentially expressed, considering a threshold of 1 for the absolute value of the log2(foldchange), and a value of 1% for the unadjusted p-value.

**Table 7.** Enrichment analysis of KEGG pathways for bacterial proteins with an absolute log_2_(fold change) less than 1 between neutrophil and monocyte fraction after Ya or Ya2 normalization strategy.

| KEGG ID | Description | Gene ratio | Background ratio | p-value | adjusted p-value |
| --- | --- | --- | --- | --- | --- |
| **Ya normalization** |  |  |  |  |  |
| ype03010 | Ribosome | 47/142 | 54/557 | 7.3E<sup>-24</sup> | 3.0E<sup>-22</sup> |
| ype01200 | Carbon metabolism | 26/142 | 65/557 | 4.5E<sup>-3</sup> | 6.3E<sup>-2</sup> |
| ype00020 | Citrate cycle | 10/142 | 17/557 | 3.1E<sup>-3</sup> | 6.3E<sup>-2</sup> |
| **Ya2 normalization** |  |  |  |  |  |
| ype03010 | Ribosome | 46/121 | 50/229 | 3.4E<sup>-11</sup> | 5.1E<sup>-10</sup> |

### Proteomic signature of *Y. pestis* grown in human plasma

To decipher bacterial adaptation to the host environment, a gene set enrichment analysis (GSEA) was performed on the gene ontology (GO) biological process terms for the 9 comparisons of *Y. pestis* growths in laboratory media and human plasma. Most of the enriched terms between samples grown in different media were related to metabolism, such as amino acid processes, purine synthetic processes such as inosine monophosphate (IMP) biosynthesis (depleted in I37 compared to M9 and plasma), tricarboxylic acid cycle (enriched in the I25 and I37 inocula), or carbohydrate transport (enriched in the I25 derived cultures) (Fig 5, Table 4). In particular, methionine biosynthesis-related proteins such as MetE (5-methyltetrahydropteroyltriglutamate--homocysteine methyltransferase), MetR (HTH-type transcriptional regulator), MetF (methylenetetrahydrofolate reductase) or MetAS (homoserine O-succinyltransferase) were enriched in plasma compared to M9 (Fig 5, 6A), consistent with adaptation to nutrient composition of plasma and in line with transcriptomic observations in murine models of plague (Sebbane et al. 2006; Lathem et al. 2005). In addition, iron transport was enriched in plasma samples compared to LBH or M9, represented by the proteins involved in biosynthesis of the yersiniabactin siderophore and other iron uptake systems such as YfeAB (Fig 5, 6B, Table 4), consistent with the low concentration of free iron in human plasma and extending previous transcriptomic data acquired in our laboratory (Chauvaux et al. 2007). We also observed higher levels of proteins encoded by the virulence plasmid pCD1, including T3SS effectors, in M9 compared to plasma and LBH (Fig 5, 6C, Table 3). Low calcium is one of the stimuli inducing T3SS expression and secretion (Perry et al. 1997). The low concentration of calcium in M9, i.e., 0.1 mM, compared to 1 mM of free ionized calcium in human plasma (Bisello et al. 2008), even in the presence of the calcium ion chelator acid-citrate-dextrose (ACD) serving as anti-coagulant, may thus be a trigger. Another signal sensed by *Y. pestis* RNA thermometers to activate expression of the T3SS regulon is the human host temperature of 37°C (Perry et al. 1997). Accordingly, when I25 and the other proteomes derived from 37°C cultures were compared, T3SS effector proteins were produced more abundantly at 37°C than 25°C (Fig 5, 6D, Table 3). Similarly, the *caf* operon encoding *Y. pestis* pseudocapsule is highly upregulated at 37°C (Demeure et al. 2019). Expectedly, we detected high levels of Caf1M, Caf1A, Caf1, and the regulator Caf1R when *Y. pestis* was grown at 37°C (I37, I25-M9, I25-Plasma, I37-M9, I37-Plasma) but not at 25°C (I25) (Table 2). In addition, pseudocapsule proteins were more abundant when cultures were started with a 37°C inoculum (I37-M9 and I37-Plasma) than with a 25°C inoculum (I25-M9 and I25-Plasma) (Table 2). Thus, *Y. pestis* growth in human plasma is characterized by a radical metabolic switch, induction of iron uptake machineries and calcium- and temperature-dependent expression of important virulence determinants.

**Figure 5.**
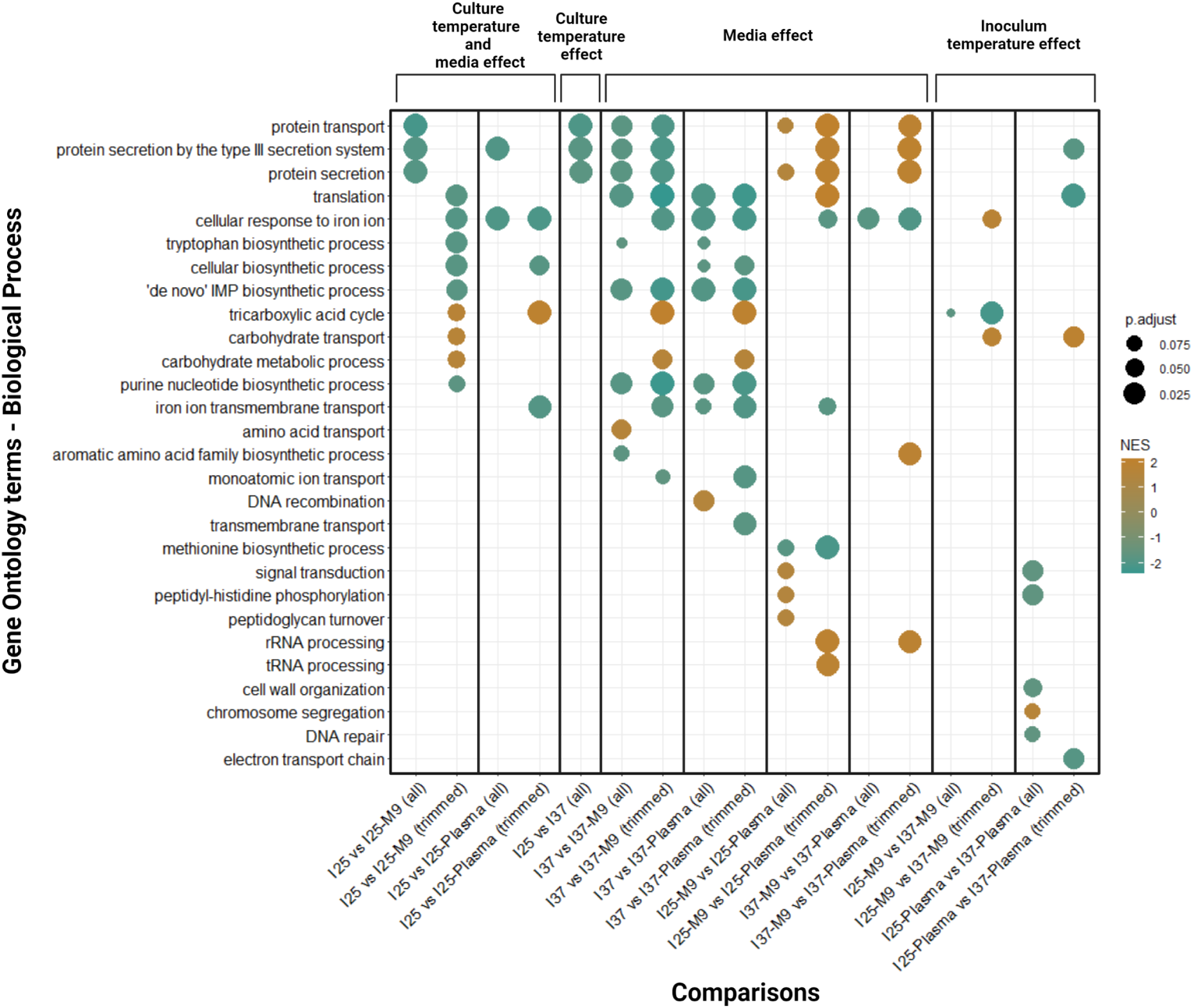
Dot plot representing the gene set enrichment analysis (GSEA) of the biological process terms from the gene ontology (GO) for the 9 proteomic comparisons. Each comparison is analyzed twice: i) with the whole data set comprising the proteins only present in one or the other conditions (marked “all”) and ii) with the proteins detected only in both conditions (marked “trimmed”). The dot color represents the normalized enrichment score (NES) of the GO term in the GSEA. The dot size represents the adjusted p-value of the enriched GO term (control of the false discovery rate by the Benjamini-Hochberg correction).

**Figure 6.**
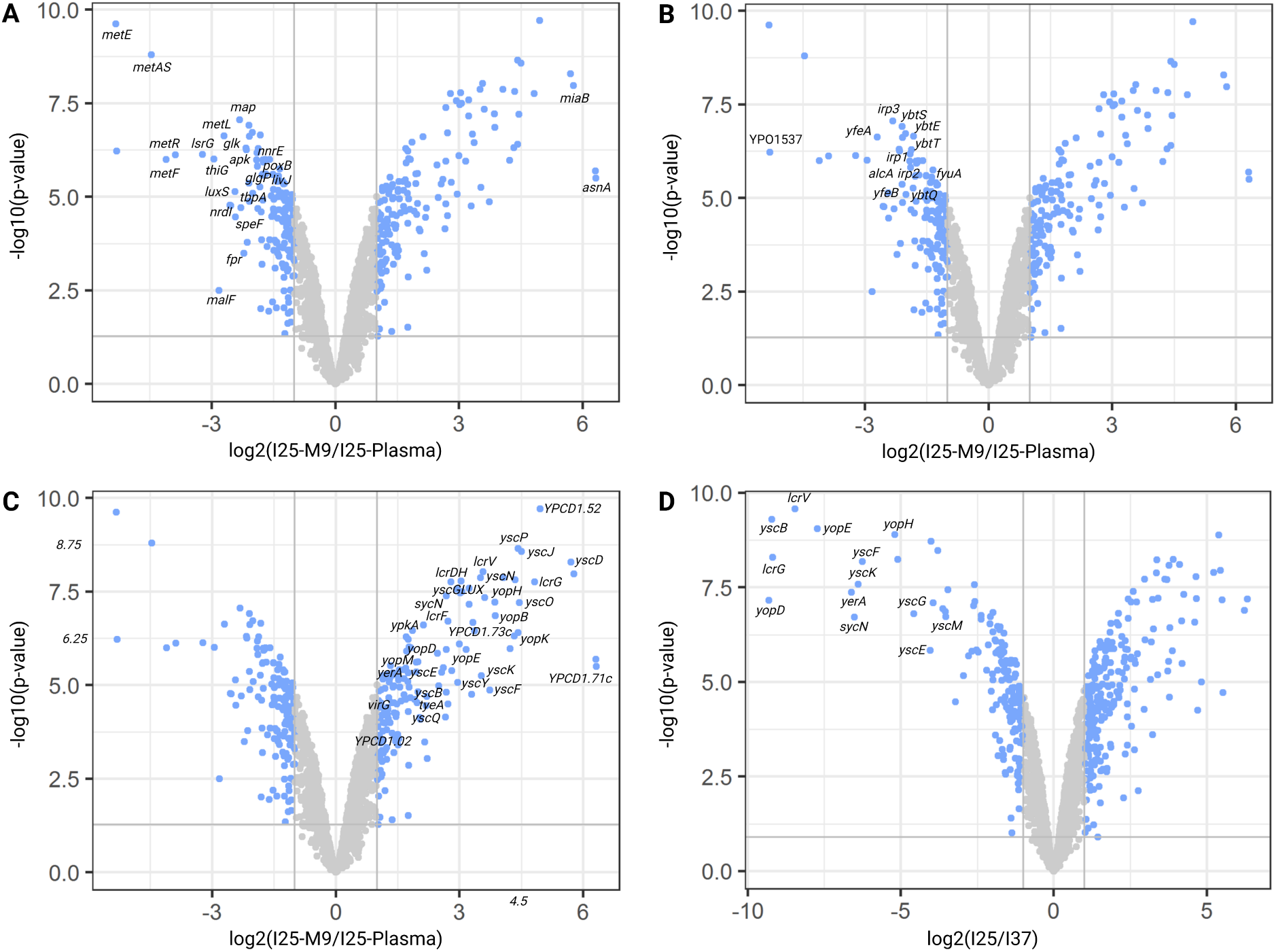
Volcano plots showing differentially abundant proteins between *Y. pestis* grown in different conditions. The displayed dots represent proteins detected in both conditions of the comparison. In blue are the proteins which are significantly differentially expressed, considering a threshold of 1 for the absolute value of the log2(foldchange), and a value of 1% for the unadjusted p-value. (A-C) M9 at 37°C with a 25°C inoculum (I25-M9) versus *Y. pestis* grown in human plasma at 37°C with a 25°C inoculum (I25-Plasma). (A) Focus on regulated metabolism proteins and the corresponding annotated genes. (B) Focus on regulated iron acquisition proteins and the corresponding annotated genes. (C) Focus on T3SS regulated proteins and the corresponding annotated genes. (D) 25°C inoculum on LBH plate (I25) versus 37°C inoculum on LBH plate (I37), with a focus on T3SS proteins and the corresponding annotated genes.

### Proteomic signature of *Y. pestis* in whole blood fractions

When analyzing the bacterial proteomes after incubation in human whole blood and immune cell purification, most of the non-secreted pCD1 encoded proteins were not differentially abundant when using the Ya2 normalization strategy. Such components include the T3S apparatus (T3SA) proteins YscL (cytoplasmic complex), YscN (ATPase), YscJ (MS-ring forming proteins), YscC (outer membrane secretin), YscW (pilotin), YscF (needle subunit), LcrV (needle tip), or regulators and chaperones such as LcrG (negative cytosolic regulator), LcrH/SycD (YopBD chaperone), YscB (YopN chaperone), YPCD1.73c (chaperone), or YerA/SycE (YopE regulator) and YscE and YscG (putative Yop translocation proteins) (Plano et al. 2013). On the other hand, secreted proteins of the pCD1 plasmid such as the T3SS effectors (here, YopH, YopE, YopJ, YopK, YpkA/YopO and YopM), the host-membrane pore-forming complex proteins YopB and YopD, the secreted gatekeeper YopN, or the secreted protein YscX (Gurung et al. 2022; Day et al. 2000) were interestingly more abundant in the monocyte fraction compared to neutrophil fraction. It was also the case for the molecular ruler YscP, serving as a needle length control protein, and the inner membrane platform protein LcrD/YscV, interacting with YscX, although both proteins were never shown to be secreted in eukaryotic cells. To validate the hypothesis that monocytes were more intoxicated with Yops than neutrophils, we used the A2 normalization strategy and validated the higher abundance of Yops in the monocyte fraction when normalized by human protein abundance (Table 8).

**Table 8.** Differential abundance of the pCD1 plasmid-encoded proteins between neutrophil and monocyte fractions, with Ya2 and A2 normalization strategies. Log_2_(fold change) less than −1 are represented in bold. T3SA: type 3 secretion apparatus.

| Locus | Protein | Role | $\log_2$(fold change)<br>Norm. Ya2 | $\log_2$(fold change)<br>Norm. A2 |
| --- | --- | --- | --- | --- |
| YPCD1.05c | YerA/SycE | regulator | -0.5 | 1.7 |
| YPCD1.06 | YopE | effector | **-3.8** | **-1.6** |
| YPCD1.19c | YopK | effector | **-2.5** | -0.3 |
| YPCD1.26c | YopM | effector | **-3.8** | **-1.6** |
| YPCD1.28c | YopD | translocon | **-4.1** | **-1.9** |
| YPCD1.29c | YopB | translocon | **-4.7** | **-2.5** |
| YPCD1.30c | LcrH/SycD | chaperone | 0.9 | 3.1 |
| YPCD1.31c | LcrV | T3SA | 0.9 | 3.1 |
| YPCD1.32c | LcrG | regulator | 0.2 | 2.4 |
| YPCD1.34c | LcrD/YscV | T3SA | **-9.3** | **-6.7** |
| YPCD1.36c | YscX | secreted | **-2.4** | -0.3 |
| YPCD1.39c | YopN | secreted regulator | **-3.3** | **-1.1** |
| YPCD1.40 | YscN | T3SA | -0.1 | 1.9 |
| YPCD1.42 | YscP | T3SA | **-9.0** | **-4.3** |
| YPCD1.48 | YscW | T3SA | **-1.5** | 0.8 |
| YPCD1.51 | YscB | chaperone | -0.3 | 2.0 |
| YPCD1.52 | YscC | T3SA | 0.4 | 2.6 |
| YPCD1.54 | YscE | T3SA | -0.4 | 1.8 |
| YPCD1.55 | YscF | T3SA | 0.3 | 2.5 |
| YPCD1.56 | YscG | T3SA | -0.8 | 1.5 |
| YPCD1.59 | YscJ | T3SA | -0.7 | 1.2 |
| YPCD1.61 | YscL | T3SA | -0.9 | 1.2 |
| YPCD1.67c | YopH | effector | **-3.1** | -0.9 |
| YPCD1.71c | YopJ | effector | **-4.5** | **-2.3** |
| YPCD1.72c | YpkA/YopO | effector | **-3.3** | **-1.1** |
| YPCD1.73c |  | chaperone | -0.4 | 1.7 |

### Human proteome signature upon *Y. pestis* infection of whole blood fractions

We could identify 4,187 human proteins in at least one replicate of the neutrophil fraction and 3,969 human proteins in at least one replicate of the monocyte fraction (Table EV4). Among the most abundant human proteins in monocytes based on PG quantities, we identified the monocyte differentiation antigen CD14, annexin A2, interleukin-1 beta (induced in non-intoxicated macrophages by YopJ-mediated inflammasome activation in neighboring intoxicated cells (Orning et al. 2018)), the galectin-1 (shown to interact with YopJ/YopP to reduce nitric oxide production in macrophages (Jofre et al. 2021)), or the gamma-interferon-inducible lysosomal thiol reductase (Table EV4). Among the most abundant human proteins in neutrophils, we identified the carcinoembryonic antigen-related cell adhesion molecule (CEACAM) 1, 6 and 8 (also respectively known as CD66a, CD66c and CD66b, specific for granulocytes and neutrophils), the protein S100-A12, S100-A8 and S100-A9 (the most abundant proteins in neutrophils (Tardif et al. 2015)), S100-P, annexin A3 (induced in neutrophils during sepsis (Toufiq et al. 2020)) or galectin-10. The most abundant human proteins in both fractions were the histone proteins (H2B type 1-O, H2A type 1-B, type 3, type 1-C, H4, H3.1, H1.3), cytoskeleton proteins (actin cytoplasmic 1 and 2, profilin), hemoglobin and albumin, but most interestingly we could find many neutrophil components such as S100-A8 and S100-A9, and granule contents such as neutrophil defensin 3, cathepsin G, neutrophil elastase, lactoferrin, azurocidin, lysozyme C or myeloperoxidase.

Using the H and H2 normalization strategies and performing a GSEA on biological process GO term, we identified neutrophil-specific terms in the most enriched processes in the neutrophil fraction, such as “neutrophil activation” or “leukocyte degranulation”, as well as terms related to actin cytosqueleton and motility, or carbohydrates metabolism (Fig 7A). On the other hand, enriched terms in monocytes were mainly related to translation and metabolic pathways linked to aerobic and mitochondrial respiration (Fig 7A). Among the proteins only identified in the neutrophil fraction, an enrichment of lipid metabolism-related proteins was observed. The proteins only identified in the monocyte fraction were again related to translation and mitochondria, as well as major histocompatibility complex class II (MHC II) (Fig 7B). These data are in line with the enrichment of the specific cell types we targeted. Indeed, in addition to cell-specific terms such as neutrophil granule or the monocyte MHCII, neutrophils are known to mainly rely on a glycolytic metabolism (Kumar et al. 2019; Injarabian, Devin, et al. 2019; Jeon et al. 2020), which could explain the enrichment of aerobic metabolism in the monocytes and a higher carbohydrate metabolism in the neutrophils.

**Figure 7.**
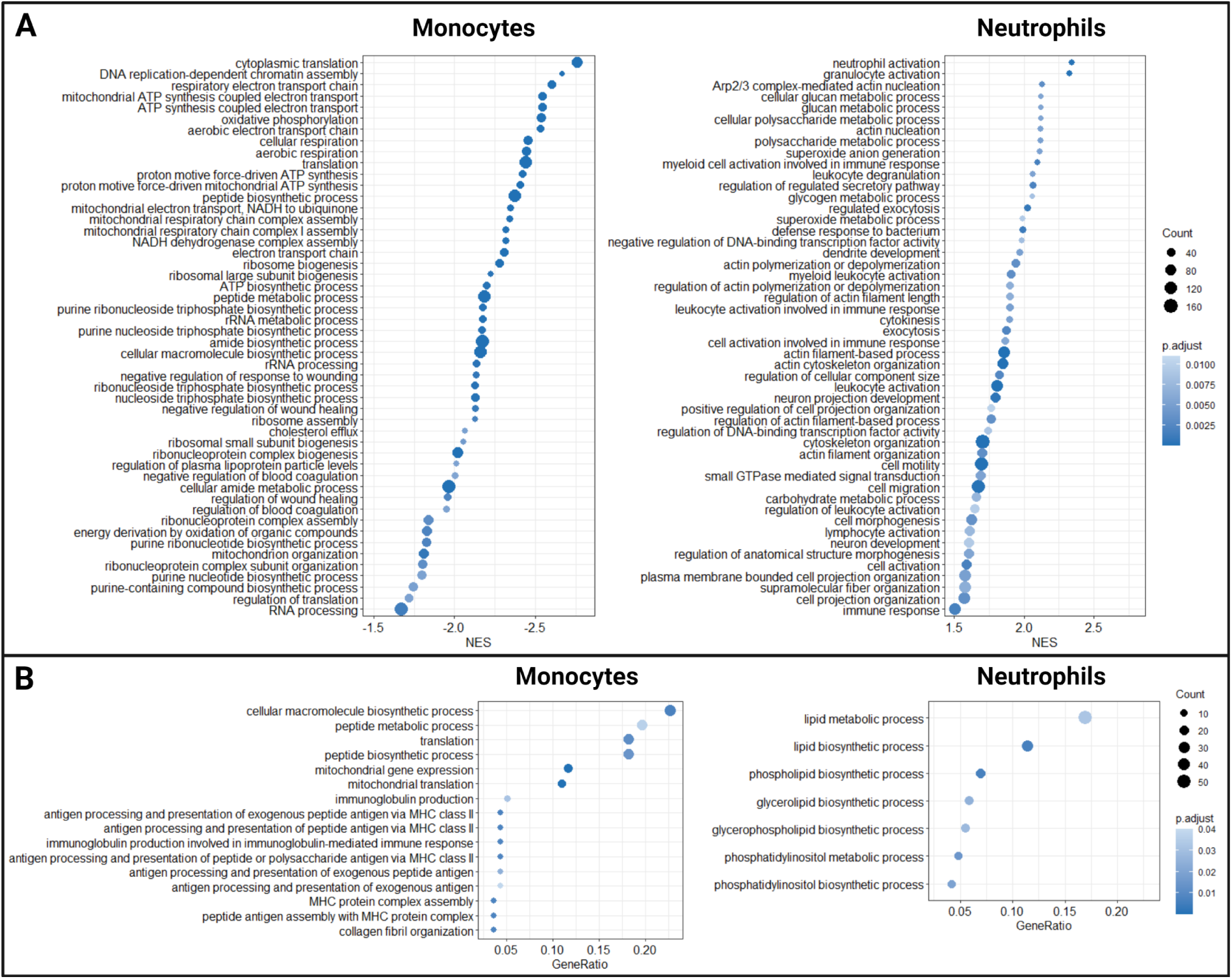
(A) Dotplots of enriched gene ontology (GO) biological processes after performing a gene set enrichment analysis (GSEA) on human protein differential abundances between the neutrophil and monocyte fractions using the H2 normalization strategy, with a focus on the 50 most enriched GO terms in monocytes (left) or in neutrophils (right). Data are ordred by normalized enrichment score (NES). (B) Dotplots of enriched GO biological processes after performing an enrichment analysis on human proteins only detected in monocytes (left) or in neutrophils (right) with the H normalization strategy. The dot size represents the number of proteins counted in this GO term. Terms are sorted by gene ratio. The dot color represents the adjusted p-value of the enriched GO term (control of the false discovery rate by the Benjamini-Hochberg correction).

## Discussion

Dual proteomics to study bacterial host-interaction in *ex vivo* or *in vivo* conditions is challenging due to the high complexity of the host proteome and the high dynamic range between bacterial and host protein abundance. Moreover, application of protocols in a BSL-3 environment are more difficult to implement and perform due to safety procedures and regulation.

Our pipeline, based on bacterial enrichment from blood, pure culture for spectral library preparation, host cell purification for sample complexity reduction, and a safe and efficient TFA-based protein extraction on minute amount of samples, could overcome all these problematics. Using a *Y. pestis* model of bacteremia, this pipeline allowed us to work in a BSL3 environment fulfilling the French regulation on MOT agents, and to perform a dual proteomic study in a highly-complex matrix, human whole blood. First, we benchmarked and validated the TFA-based protein extraction SPEED method in a BSL-3 environment, confirming bacterial inactivation upon protein extraction, and allowing comparison of bacterial protein abundances after incubation in laboratory media or human plasma. We validated and extended previous transcriptomic data (Chauvaux et al. 2007) by obtaining more sensitive and functionally relevant data. We then developed a new and affordable pipeline based on the SPEED method to study bacteria incubated in human whole blood and identified more than 5,500 proteins in the cellular fractions, composed of around 4/5 of human proteins and 1/5 of bacterial proteins, validating the possibility to use this pipeline for dual proteome studies. Cellular enrichment was performed under a hood in a BSL-3 environment before bacterial inactivation upon protein extraction, eliminating the need for expensive cell-sorting instrument or fixation step. Differential analysis of bacterial protein abundances showed a higher injection of *Y. pestis* T3SS effectors in monocytes than in neutrophils, suggesting a preferential targeting of monocytes, or a higher survival of bacteria in contact with monocytes compared to neutrophils. The normalization method was validated by the fact that most of the translation-associated proteins such as the ribosomal proteins, and metabolic pathway proteins, were not differentially abundant between both fractions. Human proteome analysis confirmed the good purification of CD14+ and CD66b+ cells, based on cell-specific GO terms and cell metabolic pathways. Strikingly, a very high abundance of neutrophil components was observed in both fractions. This could suggest degranulation of neutrophils and phagocytosis or macropinocytosis of granules and/or dead neutrophils by monocytes. Furthermore, it was recently shown that A100A8/A9 were not released along with degranulation but upon formation of neutrophil extracellular traps (NETs), suggesting activation of NETosis in our infection model (Sprenkeler et al. 2022), as S100-A8 and A9 were not only found abundantly in neutrophils but also in monocytes.

Based on our methodology, future studies could consist in comparing fraction proteomes of infected and non-infected blood samples, to better understand the immune response to infection in a relevant *ex vivo* environment, and to expand fractionation to other cell types such as platelets, T-cells or red blood cells, depending on the relevance for the pathogen under study. Our pipeline, using a small amount of blood, could be applied to blood of bacteremic animals in *in vivo* conditions. Further improvement in sensitivity could be achieved by implementing hybrid spectral libraires based on DDA and DIA runs (Willems et al. 2021), as well as using the sensitive bacterial proteome first obtained in human plasma to enhance the spectral library used in whole blood.

Challenges remain to better characterize bacteria in the bacteremic phase of diseases. One of them is the comparison of the bacterial proteome between high-complexity and low-complexity samples. We could indeed compare bacterial protein abundance between bacteria associated to neutrophils and monocytes as the host proteome background was very similar. Similarly, we could compare the proteomes of bacteria incubated in LB or human plasma, as a simple centrifugation and successive washes could separate bacteria from plasma. However, it remains difficult to compare protein abundance of cell-associated bacteria with that of a pure bacterial culture in laboratory media. One solution could be to add a host protein background to the bacterial pure culture during protein extraction to mimic sample complexity without altering protein expression. Another challenge is the study of free bacteria, as our technique relies on cell enrichment. A way to solve this issue could be to use bacteria-specific antibodies coupled to magnetic beads, such as antibodies against the F1 pseudocapsule for *Y. pestis*, and enrich bacteria from the blood. However, this approach would presumably also enrich cell-associated bacteria, and a second orthogonal separation method such as differential centrifugation would be required to separate cell-free and cell-associated bacteria. Moreover, the short life span of neutrophils once blood is drawn from donors at atmospheric level of oxygen, greatly limits truly relevant models of bacteremia (Monceaux et al. 2016; Injarabian, Skerniskyte, et al. 2021).

We believe that our methodological development successfully applied in challenging environments, such as BSL-3 laboratories, validates the proof of concept of studying by proteomics host-pathogen interactions of highly pathogenic bacteria in complex human samples.

## Methods

### Strains, culture media and human material

The avirulent *Y. pestis* CO92 pCD1-strain was used for protein extraction setup in BSL2 environment. The fully virulent *Y. pestis* CO92 was used for validation in a BSL3 environment, and incubation in human plasma and whole blood. Bacteria were routinely grown on LB agar plates (BD Difco, reference 244520) supplemented with 0.002% pork haemin (Acros Organics, reference 345960010) (LBH plate), or in LB (BD Difco, reference 244620) under agitation at 180 rpm, at 25°C or 37°C. M9 was prepared with 47.8 mM Na_2_HPO_4_ (Sigma, reference S7907), 22 mM KH_2_PO_4_ (Sigma, reference P0662), 8.6 mM NaCl (Sigma, reference 31434), 18.7 mM NH_4_Cl (Sigma, reference 09718), 2 mM MgSO_4_, 0.1 mM CaCl_2_, 1% glucose, 1% casamino acid and 1 mM thiamine-HCl. Frozen human plasma and fresh human blood anticoagulated with citrate dextrose were provided by the Etablissement Français du Sang (EFS). Whole blood was inoculated with a 25°C preculture of *Y. pestis* grown on LBH plate. For bacterial enumeration, bacteria were serially diluted in phosphate-buffered saline (PBS) and plated on LBH agar plates, which were incubated at 28°C for 48 h. When enumerating bacteria after incubation in human whole blood, 0.1% triton X100 was added in the first dilution to lyse human cells. Viable purified cells were counted in a Malassez chamber after addition of 0.1% Trypan blue at a 1:1 ratio.

### Urea-based protein extraction

After culture, bacteria were centrifuged and the pellet was washed 3 times with cold PBS. Bacteria were resuspended in 1 mL of urea lysis buffer composed of 8M urea, 100 mM TrisHCl pH 8.5, vortexed, transferred in a 2 mL tube with 0.1 mm glass beads (Micro-organism lysing VK01, Bertin Corp), and lysed in a Precellys24 (Bertin Corp) at 4°C for 90 seconds at 5,500 rpm. Lysates were centrifugated 5 minutes at 15,000 *g* and 500 µL of supernatant were concentrated using an Amicon^®^ Ultra 30 kDa filter (Merck Millipore) following centrifugation at 14,000 *g* for 20 minutes. Proteins were resuspended in 450 µL 8M urea, 100mM TrisHCl pH 8.5, before addition of 50 µL of tris(2-carboxyethyl)phosphine (TCEP) 100 mM and 2-chloroacetamide (CAA) 400 mM and incubated for 5 min at 95°C. Samples were centrifugated at 14,000 *g* for 10 minutes, and 300 µL of ammonium bicarbonate (ABC) 100 mM pH 8.0 were added. Following centrifugation 3 times at 14, 000 *g* for 10 minutes, proteins were resuspended in 350 µL ABC 100 mM and quantified as explained below.

### SDS-based protein extraction

After culture, bacteria were centrifuged and the pellet was washed 3 times with cold PBS. Bacteria were resuspended in 1 mL of a SDS lysis buffer composed of 4% SDS, 100mM TrisHCl, 10 mM TCEP, 40 mM CAA, incubated for 5 minutes at 95°C and sonicated for 10 seconds. Samples were centrifuged at 16,000 *g* for 5 minutes, supernatants were concentrated with an Amicon^®^ Ultra 30 kDa filter (Merck Millipore) following centrifugation at 14,000 *g* for 20 minutes. Proteins were resuspended in 450 µL 8M urea, 100 mM TrisHCl pH 8.5, followed by centrifugation 2 times at 14,000 *g* for 20 minutes. 100µL 8M urea, 100 mM TrisHCl pH 8.5 were added, followed by centrifugation 3 times at 14,000 *g* for 15 minutes. Then 300µL ABC were added, followed by centrifugation 3 times at 14,000 *g* for 10 minutes. Proteins were resuspended in 350 µL ABC 100 mM, pH 8.0 and quantified as explained below.

### TFA-based protein extraction

After culture, bacteria were centrifuged and the pellet was washed 3 times with cold PBS. Bacteria were resuspended in 5 volumes of TFA compared to the pellet (20 µl of TFA for a pellet of 4 µL) incubated for 10 minutes and transferred to a sterile 1.5 mL protein low-binding tube. 10 volumes, compared to the TFA volume (200 µL) of 2M Tris were added, then 24 µL of TCEP 100mM, CAA 400 mM (for a final concentration of TCEP at 10 mM and CAA at 40 mM). Samples were incubated at 95°C for 5 minutes and proteins were quantified as explained below. Volumes were adjusted to achieve the desired quantity of proteins to digest, and 5 volumes of water were added.

### Protein quantification

Bradford, BCA and Qubit assays were performed as described by the supplier. Bovine serum albumin (BSA) standard resuspended in 8M urea, 100 mM TrisHCl pH 8.5 or 4% SDS, 100 mM TrisHCl pH 8.5 or 10% TFA-2M Tris was used to assess buffer compatibility with the different assays. Tryptophan standard was performed using pure L-tryptophan dissolved in distilled water. 50 µL of samples were aliquoted in black 96-well (or alternatively 384-well) plates and measured with a Xenius spectrophotometer (or alternatively Synergy H1M microplate reader - BioTek) with an excitation wavelength of 280 nm and an emission wavelength of 360 nm.

### Protein digestion and desalting

For the comparison of urea-based, SDS-based and TFA-based protein extractions, proteins were digested using Sequencing Grade Modified Trypsin (Promega - V5111) with a 1:50 ratio (enzyme:protein) at 37°C for 12 h before stopping the digestion by addition of TFA to reach a final pH lower than 2. Digested peptides were desalted on 50 mg Sep-Pak C18 cartridge (Waters - WAT054955). Peptides were eluted twice with a acetonitrile (ACN) 50%, formic acid (FA) 0.1% buffer and once with an ACN 80%, FA 0.1% buffer. Finally, the peptide solutions were speed-vac dried and resuspended in ACN 2%, FA 0.1% buffer. For each sample, absorbance at 280 nm was measured with a Nanodrop^TM^ 2000 spectrophotometer (Thermo Scientific) to inject an equivalent of DO = 1. For *Y. pestis* incubation in human plasma and in human whole blood, digested peptides were desalted using the Stage-Tips method (Rappsilber et al. 2007) using C18 Empore disc and eluted with ACN 80%, FA 0.1%. Finally, the peptide solutions were speed-vac dried and resuspended in ACN 2%, FA 0.1% buffer. For each sample, absorbance at 280 nm was measured with a Nanodrop^TM^ 2000 spectrophotometer (Thermo Scientific) to inject an equivalent of DO = 1.

### Peptide mixing and pre-fractionation

A pool of 100 µg of all SPEED digested samples was used to obtain a spectral library for DIA. To do this, equivalent amounts of each digested sample were pooled before proceeding to a peptide fractionation using a manual workflow based on Stage-Tips or an automatic workflow with the AssayMAP Bravo (Agilent).

Manual workflow was performed with a poly(styrene-divinylbenzene) reverse phase sulfonate (SDB-RPS) Stage-Tips method as described in (Rappsilber et al. 2007; Kulak et al. 2014). The pooled sample (20 µg) was loaded into 3 SDB-RPS (Empore^TM^, 66886-U) discs stacked on a P200 tip and 8 serial elutions were applied as following: elution 1 (Ammonium formate (AmF) 60 mM, ACN 20%, FA 0.5%), elution 2 (AmF 80 mM, ACN 30%, FA 0.5%), elution 3 (AmF 95 mM, ACN 40%, FA 0.5%), elution 4 (AmF 110 mM, ACN 50%, FA 0.5%), elution 5 (AmF 130 mM, ACN 60%, FA 0.5%), elution 6 (AmF 150 mM, ACN 70%, FA 0.5%) and elution 7 (ACN 80%, ammonium hydroxide 5%).

Automatic workflow was performed using the AssayMAP Bravo with the Fractionation v1.1 protocol. The dried pooled sample (80 µg) was resuspended in 20 mM ammonium formate, pH 10 before a high pH reverse phase fractionation. RPS cartridge (Agilent Technologies, 5 µL bead volume, G5496-60033) were primed with 100 µL ACN 80%, FA 0.1% and equilibrated with 70 µL 20 mM AmF, pH 10. Samples were loaded at 5 µL/min followed by an internal cartridge wash and cup wash with 50 µL of 20 mM AmF, pH 10 at 5 µL/min. Step elution was performed with 60 µL of ACN 10%, 20%, 30%, 40%, 50%, and 80% in 20 mM AmF, pH 10 at 5 µL/min. A preexisting volume of 20 µL containing the same elution buffer was present in the collection plates upon elution.

All fractions were speed-vac dried and resuspended with ACN 2%, FA 0.1% before injection. For all fractions, iRT peptides were spiked as recommended by Biognosys (Biognosys - Ki-3002-1).

### LC-MS/MS for DDA and spectral libraries creation

In all proteomic analyses, a nanochromatographic system (Proxeon EASY-nLC 1200 - Thermo Scientific) was coupled online with a Q Exactive^TM^ HF mass spectrometer (Thermo Scientific).

For the comparison of urea-based, SDS-based and TFA-based protein extractions, 1 µg of peptides was injected into a reverse phase column (EASY-Spray^TM^ - ES902 - Thermo Scientific: 25cm x 75 µm ID, 2.0 µm particles, 100 °A pore size,) after an equilibration step in 100% solvent A (H_2_O, FA 0.1%).

For comparisons of *Y. pestis* protein abundances after growth in human plasma or laboratory media, 1 µg of peptides was injected into a reverse phase column (home-made column, 45 cm x 75 µm ID, 1.9 µm particles, 100 °A pore size, ReproSil-Pur Basic C18 - Dr. Maisch GmbH, Ammerbuch-Entringen, Germany) after an equilibration step in 100% solvent A (H_2_O, 0.1% FA). For creation of the spectral library, an equivalent volume of each fraction was injected into the same reverse phase column.

Peptides were eluted with a multi-step gradient from 2 to 7% buffer B (ACN 80% / FA 0.1%) in 5 min, 7 to 23% buffer B in 70 min, 23 to 45% buffer B in 30 min and 45 to 95% buffer B in 5 min at a flow rate of 250 nL/min for up to 132 min. Column temperature was set to 60°C. Mass spectra were acquired using Xcalibur software using a data-dependent Top 10 method with a survey scans (300-1700 m/z) at a resolution of 60,000 and MS/MS scans (fixed first mass 100 m/z) at a resolution of 15,000. The AGC target and maximum injection time for the survey scans and the MS/MS scans were set to 3.0×10^6^, 100 ms and 1.0×10^5^, 45 ms respectively. The isolation window was set to 1.6 m/z and normalized collision energy fixed to 28 for HCD fragmentation. We used a minimum AGC target of 2.0×10^3^ for an intensity threshold of 4.4×10^4^. Unassigned precursor ion charge states as well as 1, 7, 8 and *>*8 charged states were rejected and peptide match was disable. Exclude isotopes was enabled and selected ions were dynamically excluded for 45 seconds.

### LC-DIA-MS for DIA

For the *Y. pestis* incubation in human blood, two libraries for DIA were constructed from i) CD14 and CD66b cell enrichments + Nanotrap^®^-enriched *Y. pestis* and ii) pure culture of *Y. pestis* in LB at 37°C. Mass spectra were acquired in DIA mode with the XCalibur software using the same nanochromatographic system coupled on-line to a Q Exactive^TM^ HF Mass Spectrometer. For each sample, 1 µg of peptides was injected into a reverse phase column (home-made column, 45 cm x 75 µm ID, 1.9 µm particles, 100 °A pore size, ReproSil-Pur Basic C18 - Dr. Maisch GmbH, Ammerbuch-Entringen, Germany) after an equilibration step in 100% solvent A (H_2_O, 0.1% FA). Peptides were eluted using the same multi-step gradient and temperature. MS data was acquired using the Xcalibur software with a scan range from 295 to 1170 m/z. The DIA method consisted in a succession of one MS scan at a resolution of 60,000 and 36 MS/MS scans of 1 Da overlapping windows (isolation window = 25 m/z) at 30,000 resolution. The AGC (Automatic Gain Control) target and maximum injection time for MS and MS/MS scans were set to 3.0×10^6^, 60 ms and 5.0×10^5^, auto respectively. The normalized collision energy was set to 28 for HCD fragmentation.

### Bioinformatic analyses

MaxQuant analyses: DDA raw files were processes using Max-Quant software version 1.6.10.43 (Cox and Mann 2008) with Andromeda search engine (Cox, Neuhauser, et al. 2011). The MS/MS spectra were searched against a UniProt *Y. pestis* database (3,909 entries the 25/04/2019). Variable modifications (methionine oxidation and N-terminal acetylation) and fixed modification (cysteine carbamidomethylation) were set for the search and trypsin with a maximum of two missed cleavages was chosen for searching. The minimum peptide length was set to 7 amino acids and the false discovery rate (FDR) for peptide and protein identification was set to 0.01. The main search peptide tolerance was set to 4.5 ppm and to 20 ppm for the MS/MS match tolerance. Second peptides was enabled to identify co-fragmentation events. The “match between runs” feature was applied for samples having the same experimental condition with a maximal retention time window of 0.7 minute. One unique peptide to the protein group was required for the protein identification. Quantification was performed using the XIC-based LFQ algorithm with the Fast LFQ mode as described in Cox, Hein, et al. 2014. Unique and razor peptides included modified peptides, with at least 2 ratio count were accepted for quantification.

Spectral libraries generation: DDA raw data were directly loaded into Pulsar and spectral library generation was done using an in-house *Y. pestis* CO92 database containing 3,991 entries, including 3,915 unique proteins from the initial CO92 Sanger sequencing (Parkhill et al. 2001), reannotated by RefSeq (Accession GCF 000009065.1 ASM906, accessed in 2020) and 76 small open frame (sORF) encoded peptides (SEP) recently identified (Cao et al. 2021) and the UniProt Homo sapiens database (20,361 entries the 03/03/2023). We applied the default BGS Factory Settings for both Pulsar Search and Library Generation. Briefly, full tryptic digestion allowing two missed cleavages, carbamidomethylation as a fixed modification on all cysteines, oxidation of methionines, and protein N-terminal acetylation as dynamic modifications were set. We used the search archive properties to create specific *Y. pestis* and human libraries as well as a combined library with a control of the FDR. Spectronaut analyses: Spectronaut v. 16.0.220606 (Biognosys AG) (Bruderer et al. 2015) was used for DIA-MS data analyses. Data extraction was performed using the default BGS Factory Settings. Briefly, for identification, both precursor and protein FDR were controlled at 1%. For quantification, peptides were grouped based on stripped sequences and Qvalue was used for precursor filtering. MaxLFQ was used with no imputation and no cross run normalization strategies.

### Statistical analysis

Protocol comparison and plasma analysis: To find the proteins more abundant in one condition than another, the LFQ intensities quantified using MaxQuant were compared. Only proteins identified with at least one peptide that was not common to other proteins in the FASTA file used for the identification (at least one “unique” peptide) were kept. Additionally, only proteins with at least two intensity values in one of the two compared conditions were kept for further statistics. Proteins absent in a condition and present in another are put aside. These proteins can directly be assumed differentially abundant between the conditions. After this filtering, intensities of the remaining proteins were first log-transformed (log2). Next, intensity values were normalized by median centering within conditions (Giai Gianetto 2023). Missing values were imputed using the impute.mle function of the R package imp4p (Giai Gianetto et al. 2020). Statistical testing was conducted using a limma t-test thanks to the R package limma (Ritchie et al. 2015). An adaptive Benjamini-Hochberg procedure was applied on the resulting p-values thanks to the function adjust.p of the cp4p R package (Giai Gianetto et al. 2016) using the robust method described in Pounds et al. 2006 to estimate the proportion of true null hypotheses among the set of statistical tests. The proteins associated to an adjusted p-value inferior to a FDR (false discovery rate) level of 1% and an absolute log2(fold-change) superior to 1 have been considered as significantly differentially abundant proteins. Finally, the proteins of interest were therefore those which emerged from this statistical analysis supplemented by those which were considered to be present from one condition and absent in another.

Whole blood analysis: To find the proteins more abundant in one condition than in another, the intensities quantified using Spectronaut were compared. Only proteins identified with at least one peptide that is not common to other proteins in the FASTA file used for the identification (at least one “unique” peptide) were kept. Depending on the normalization and differential analysis strategy used, only proteins with at least one intensity values in one of the two compared conditions, or at least two intensity values in both compared conditions were kept for further statistics. In the first case, proteins absent in a condition and present in another are put aside. These proteins can directly be assumed differentially abundant between the conditions. After this filtering, intensities of the remaining proteins were first log-transformed (log2). Next, depending on the normalization strategy, intensity values only for bacterial or human proteins were normalized either by median centering within conditions or by median centering on all conditions (Giai Gianetto 2023). This method consists in estimating the median of intensities in a sample j (*med_j_*), to substract this median to all the intensities *x* in the sample j and to add the average of median intensities (*y* = *x − med_j_* + *average*(*med_j_*)). Missing values were imputed using the impute.slsa function of the R package imp4p (Giai Gianetto et al. 2020). Statistical testing was conducted using a limma t-test thanks to the R package limma (Ritchie et al. 2015). An adaptive Benjamini-Hochberg procedure was applied on the resulting p-values thanks to the function adjust.p of the cp4p R package (Giai Gianetto et al. 2016) using the robust method described in Pounds et al. 2006 to estimate the proportion of true null hypotheses among the set of statistical tests. The proteins associated to an adjusted p-value inferior to a FDR (false discovery rate) level of 1% and an absolute log2(fold-change) superior to 1 have been considered as significantly differentially abundant proteins. Finally, the proteins of interest were therefore those which emerged from this statistical analysis supplemented by those which were considered to be present from one condition and absent in another.

We used the R package clusterProfiler v4.6.2 for enrichment and gene set enrichment analysis (Wu et al. 2021). For the bacterial proteome in plasma, the GSEA function was used based on UniProt accession number and mapped to an in-house annotation file extracted from the CO92 Gene Ontology terms downloaded from the QuickGo website accessed in April 2023 (Binns et al. 2009). The enrichKEGG function was used with the online KEGG dataset for the ype organism accessed in April 2023, based on the CO92 loci. Analysis were performed with the detected proteins as statistical background. For the human proteome analysis, the gseGo was used with the org.Hs.eg.db v3.16.0 as annotation database and statistical background, and the gseKEGG was used with the online annotation for the hsa organism accessed in April 2023 as annotation database and statistical background.

### Comparison of bacterial proteome after growth in human plasma or laboratory media

The fully virulent CO92 strain was cultivated overnight for 20 h on LBH plate at 25°C or 37°C. Frozen human plasma (Donor A) was thawed, centrifuged at 8,600 *g* for 10 minutes and filtered on 0.22 µm filter to remove precipitates. Bacteria were resuspended in 3 mL PBS and the optical density at 600 nm (OD600) was adjusted to 0.25 for the 25°C preculture and 0.67 for the 37°C preculture. 2 mL of preheated human plasma or 2 mL of preheated M9 media were inoculated with 200 µL of each suspension and incubated at 37°C for 8 h under agitation at 150 rpm in 14 mL polypropylene tubes with round bottom, in 3 replicates. Bacteria grown for 8 hours and the inocula were pelleted by centrifugation and washes 3 times with cold PBS, followed by centrifugation at 10,000 *g* for 10 min at 4°C. Bacteria were then lysed with 10 µL TFA for the inocula and bacteria grown in M9, and 20 µL TFA for bacteria grown in plasma and processed as explained above in “TFA-based protein extraction”. To assess reproducibility of protein expression between different plasma donors, the same experiment was performed with donor A and 3 other donors (B, C and D).

### Comparison of monocyte- and neutrophil-associated bacterial proteome

The fully virulent CO92 strain was grown overnight for 20 h on LBH plate at 25°C. Bacteria were resuspended in 3 mL PBS and the OD600 was adjusted to 0.25. 200 µL of the inoculum were added to 2 mL of preheated human blood at 37°C, and 2 mL of preheated LB in 14 mL polypropylene tubes with round bottom, in 5 replicates. Suspensions were incubated at 37°C for 8 h under agitation at 180 rpm.

For immune cell purifications, 1 mL and 3 mL of the blood cultures were aliquoted, filtered on 30 µm mesh to remove clogs and aggregates, in 5 replicates. StraightFrom^®^ Whole Blood kits (Miltenyi Biotec) were used to selectively enrich either monocytes (CD14 beads) or neutrophils (CD66b beads). 50 µL of CD66b beads were added to 1 mL of blood culture and 150 µL CD14 beads were added to 3 mL of blood culture, incubated for 15 minutes at 4°C and transfered in conditioned Whole Blood Columns (Miltenyi Biotec) following the manufacturer’s instructions. After 2 washes with PBS, 2 mM EDTA, 0.5% BSA, cells were eluted with 4 mL Whole Blood Column Elution Buffer (Miltenyi Biotec) and centrifuged at 500 *g* for 10 minutes at 4°C. The supernatants were removed, cells and associated bacteria were lysed with 80 µL TFA and proteins were processed as explained above in “TFA-based protein extraction”.

For the first library construction, bacteria enriched from whole blood with magnetic Nanotrap^®^ Microbiome A particles (Ceres Nanosciences, Inc) after the 8 h-incubation, in 5 replicates. In 15 mL tubes, 1 mL of Nanotrap^®^ beads were added to 4 mL of blood culture, briefly vortexed, and incubated at room temperature for 30 min with regular quick vortexing. After the addition of 6 mL of PBS, samples were vortexed and put on a magnetic rack for 10 minutes. Supernatants were discarded, the tubes were removed from the magnetic rack, 1 mL of PBS was added and samples were vortexed. The mix of beads and cells were then transferred to a sterile 1.5mL protein low-binding tube and pelleted on a magnetic rack for 10 minutes. The supernatant was discarded, and the pellet was washed a second time with 1 mL PBS. After a third magnetic pelleting, the cell pellet was lysed with 100 µL TFA and processed as explained above in “TFA-based protein extraction”.

For the second library construction, 4 mL of the 8 h-incubation in LB were pelleted by centrifugation at 4,000 *g* for 15 minutes at 4°C and washed 3 times with 4 mL cold PBS, in 5 replicates. The pellet was then lysed with 80 µL TFA and processed as explained above in “TFA-based protein extraction”.

## Data availability

The mass spectrometry proteomics data have been deposited to the ProteomeXchange Consortium via the PRIDE partner repository (Perez-Riverol et al. 2022) with the dataset identifier PXD042837.

## Acknowledgments

We are grateful to all members of the *Yersinia* research unit and the *Yersinia* National Reference Laboratory, WHO Collaborating Research and Reference Centre for Plague FRA-140 for insightful discussions.

The project received funding from Institut Pasteur, Université Paris Cité, CNRS, LabEX Integrative Biology of Emerging Infectious Diseases (ANR-10-LBX-62-IBEID), Agence de l’Innovation de Défense (AID - DGA), Fondation pour la Recherche Médicale (FDT20220401-5222) and the Inception program (Investissement d’Avenir grant ANR-16-CONV-0005). The funders had no role in study design, data collection and interpretation, or the decision to submit the work for publication.

Most of the figures were created with BioRender.

We declare no conflict of interest.

## Author contributions

**Pierre Lê-Bury**: conceptualization; investigation; formal analysis; visualization; writing – original draft; **Thibaut Douché**: investigation; formal analysis; visualization; **Quentin Giai Gianetto**: investigation; formal analysis; visualization; **Mariette Matondo**: formal analysis; writing – original draft; funding acquisition; **Javier Pizarro-Cerdá**: formal analysis; writing – original draft; funding acquisition; **Olivier Dussurget**: conceptualization; formal analysis; writing – original draft; funding acquisition. All the authors contributed to editing the original manuscript before submission.

## Expanded View Figures

**Expanded View Figure 1.**
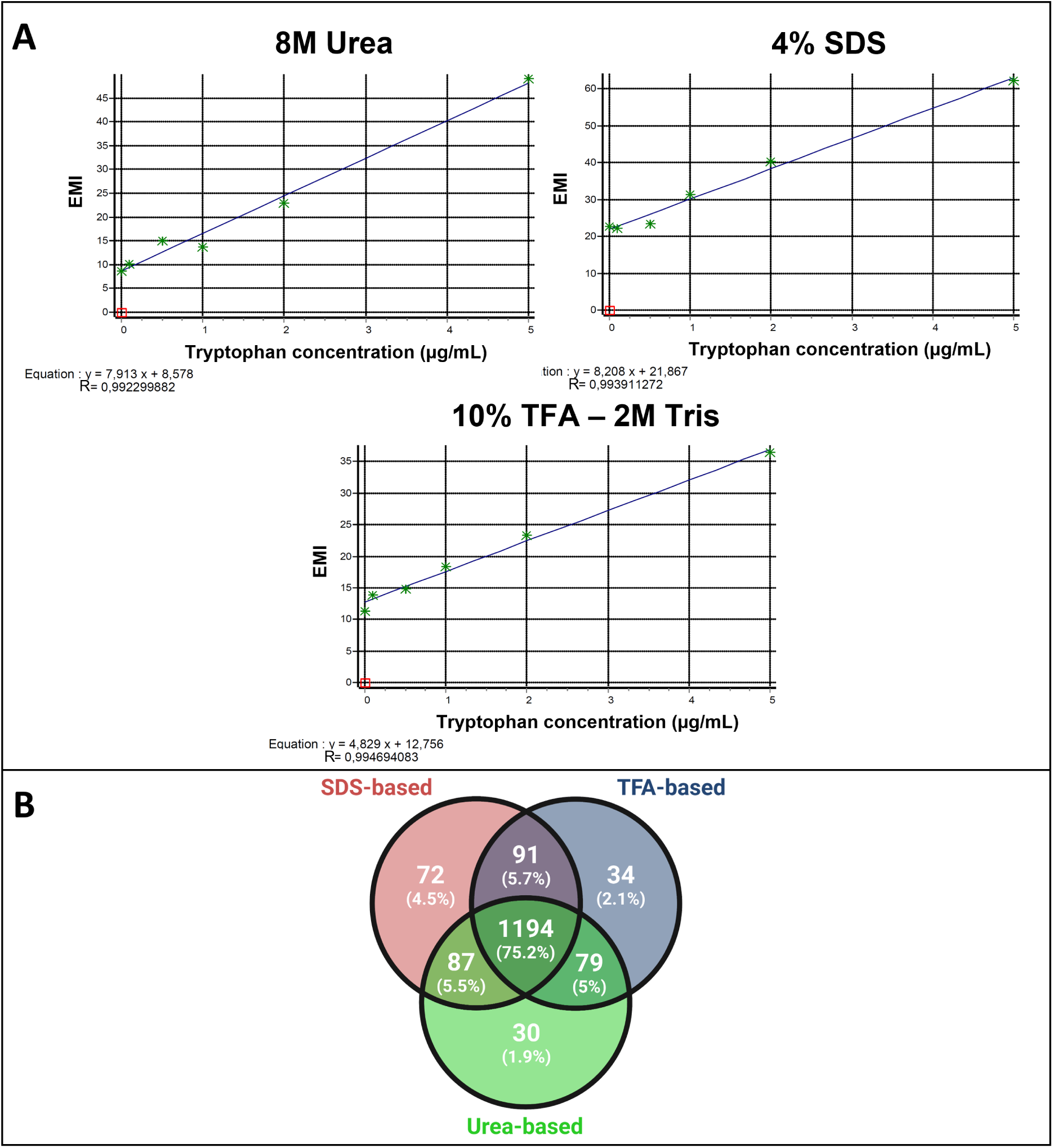
(A) Tryptophan standard measured by fluorescence (excitation: 280nm, emission: 360nm) in a plate reader for the three lysis buffers: 8M urea, 4% SDS and 10% TFA - 2M Tris. (B) Venn diagram of identified proteins repartition between the urea-based, SDS-based and TFA-based protocols.

**Expanded View Figure 2.**
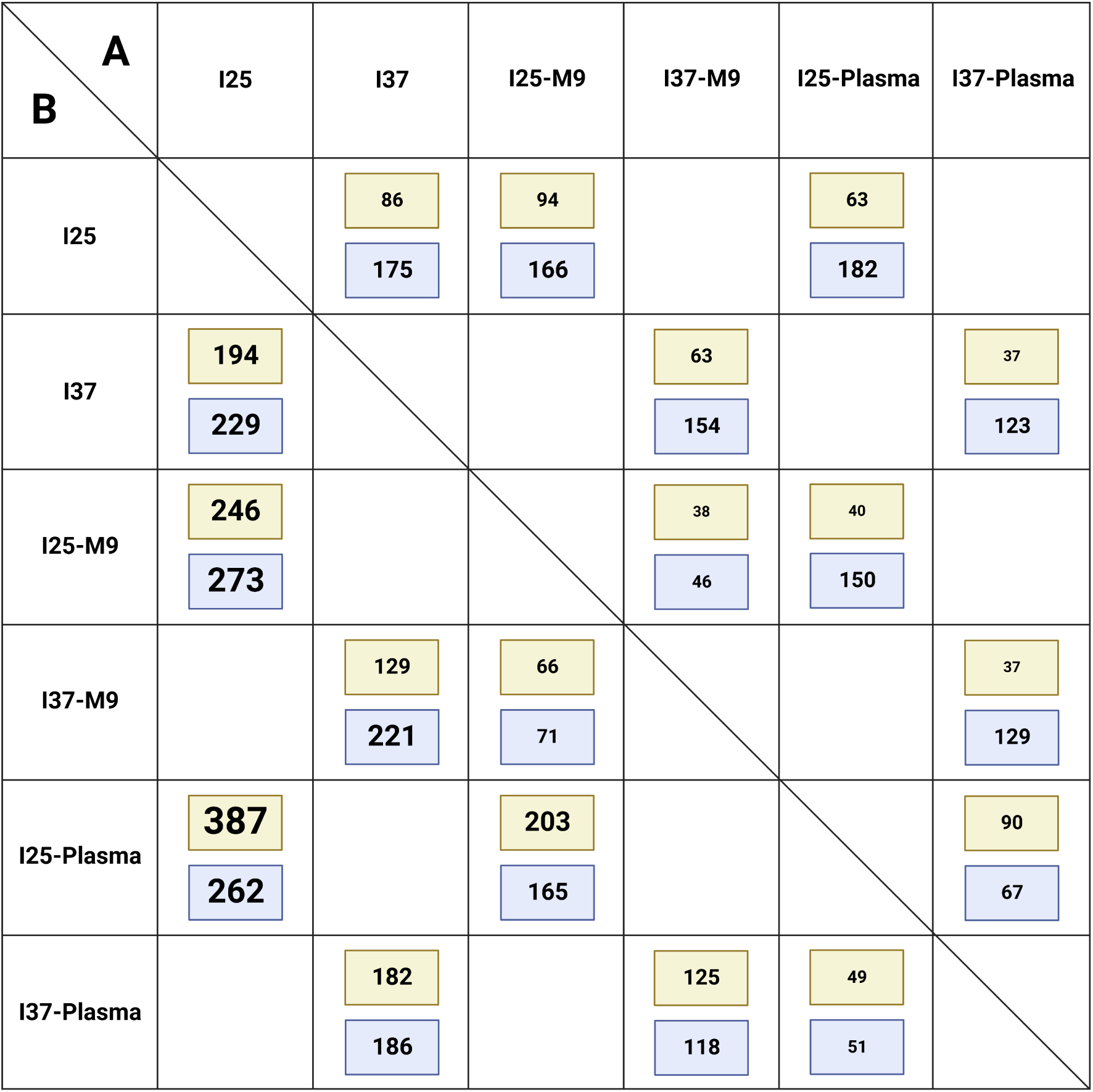
Differentially abundant proteins and proteins present and absent in the different comparisons. Each cell corresponds to the comparison A versus B (column versus row). In green are the number of proteins present in A and absent in B, in blue are the numbers of proteins present in both conditions and more abundant in A versus B.

**Expanded View Figure 3.**
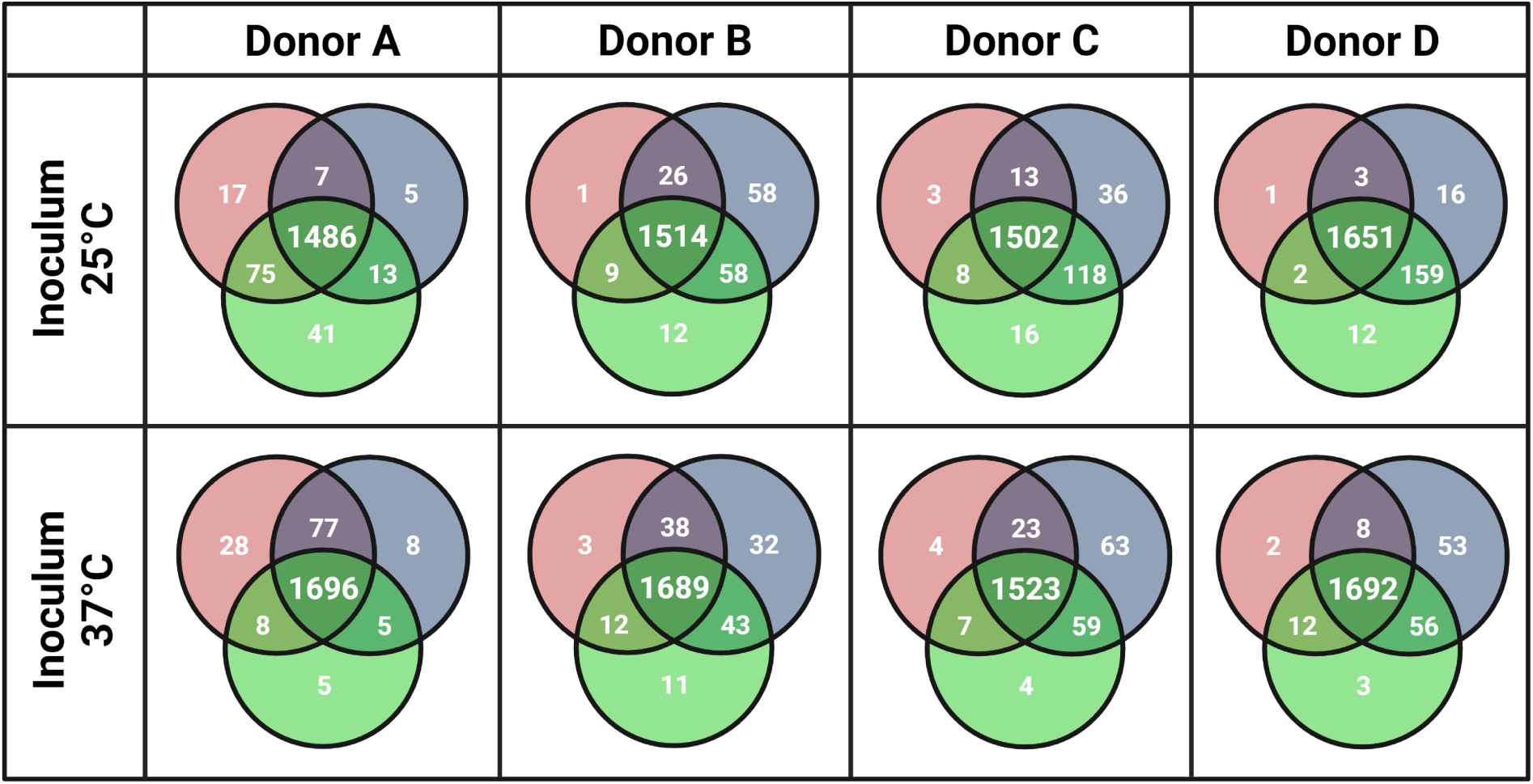
Venn diagram of number of detected proteins in replicates after *Y. pestis* growth in plasma of different donors.

**Expanded View Figure 4.**
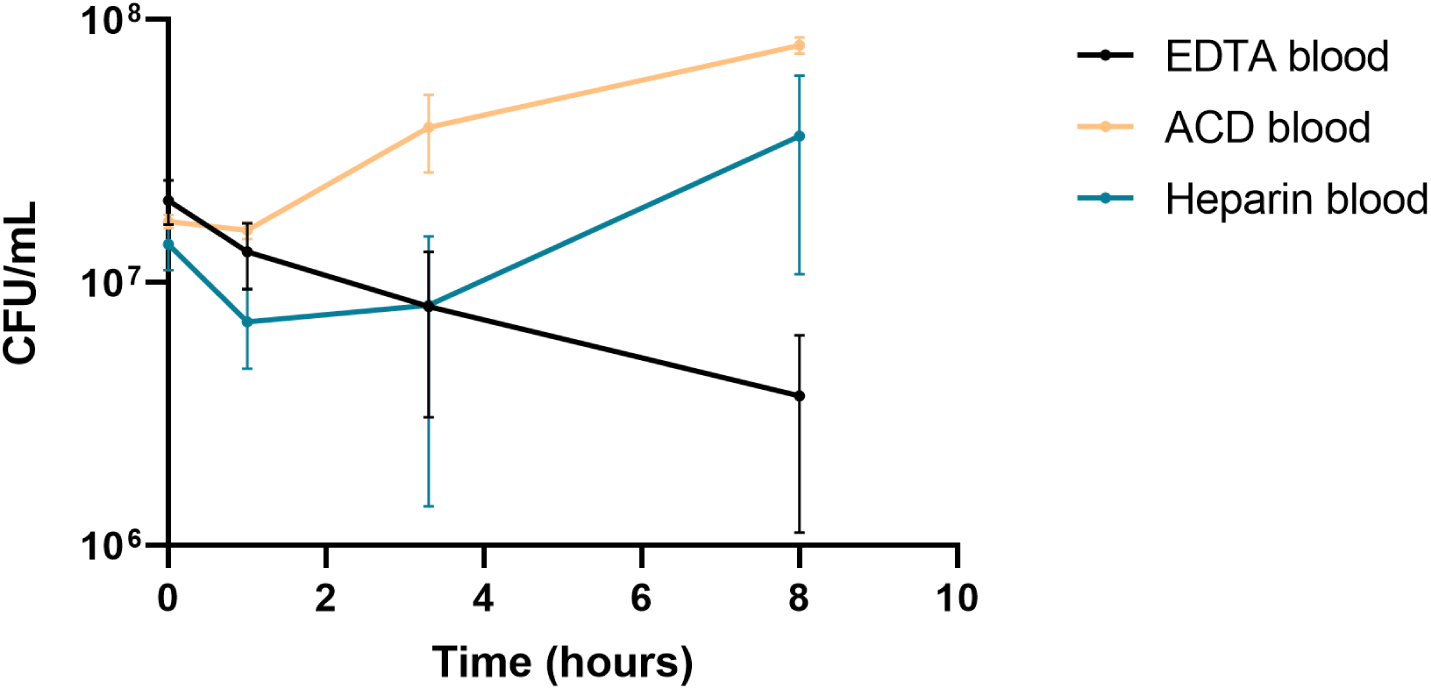
Growth of *Y. pestis* CO92 in human blood anticoagulated with EDTA, ACD or heparin.

